# Asymmetrical modulation of fear expression via GABA_B_ receptors in the mouse medial habenula

**DOI:** 10.64898/2025.12.02.691389

**Authors:** Cihan Önal, Peter Koppensteiner, Mary Muhia, Elodie Le Monnier, Ryuichi Shigemoto

**Affiliations:** Institute of Science and Technology Austria, Klosterneuburg 3400, Austria

## Abstract

The medial habenula (MHb) is implicated in regulating emotional responses to aversive events. Studies in zebrafish have identified a remarkable morphological left-right asymmetry in the dorsal habenula (zebrafish equivalent of mammalian MHb)-interpeduncular nucleus (IPN) pathway and its left-sided-specific role in modulating fear responses. However, there is little evidence for structural or functional lateralization in the mammalian MHb-IPN pathway. Here, we investigated the synaptic properties of left- and right-MHb afferents to the IPN and their roles in the expression of conditioned fear in mice. We found that each IPN neuron receives inputs from both left and right MHb, but the left MHb-originating synapses exhibit lower release probability and higher γ-aminobutyric acid type B receptor (GABA_B_R)-mediated potentiation compared to the right MHb-originating synapses. Interestingly, these asymmetrical properties persist in the inversus visceral mutant mice with normal internal organ laterality (situs solitus), but nearly disappear in those with reversed internal organ laterality (situs inversus). Behaviorally, chemogenetic inhibition of cholinergic neurons and conditional deletion of GABA_B_R in the left, but not the right, MHb significantly attenuated cue-dependent fear recall. Our results demonstrate functional asymmetry of the MHb under partial influence of the nodal flow in mice, revealing a predominant role of GABA_B_R-mediated signaling in the left MHb-IPN pathway in modulating fear memories. These findings suggest that lateralized MHb pathways could represent a fundamental principle in the neural regulation of emotion across species, but that they develop differently in zebrafish and mice.

## Introduction

Fear learning serves an adaptive purpose by aiding the organism in avoiding danger and promoting survival (LeDoux, 2000; Maren and Quirk, 2004). On the other hand, maladaptive expression of fear, especially in the absence of danger, represents a key feature in fear-based disorders such as phobias and post-traumatic stress disorder (Ressler and Mayberg, 2007; Pitman *et al*., 2012; Craske and Stein, 2016). Human and animal studies show that the acquisition, recall, and extinction of fear rely on the integrity of a triad of brain regions comprising the hippocampus, amygdala, and prefrontal cortex (Maren, 2001; Hobin, Goosens, and Maren, 2003; Shin, Rauch). Beyond the core circuitry, extensive evidence has demonstrated a key modulatory role for the habenula in classical aversive conditioning and other negative emotional states (Agetsuma *et al*., 2010; Hikosaka, 2010; Okamoto, Agetsuma, and Aizawa, 2012; Görlich *et al*., 2013; Viswanath *et al*., 2013; McLaughlin, Dani, and De Biasi, 2017; Fore *et al*., 2018; Lee *et al*., 2019).

The habenula complex is a phylogenetically conserved bilateral structure in the epithalamus that serves as a node linking the forebrain and midbrain regions (Hikosaka, 2010; Ables, Park, and Ibañez-Tallon, 2023). In zebrafish, the habenula comprises two structurally and functionally distinct sub-nuclei: the ventral and dorsal habenula (dHb), which correspond to the lateral habenula (LHb) and medial habenula (MHb) in mammals, respectively (Namboodiri, Rodriguez-Romaguera, and Stuber, 2016; Ables, Park, and Ibañez-Tallon, 2023). The LHb and MHb differ in gene expression profiles and neuroanatomical connectivity (Kim and Chang, 2005; Aizawa *et al*., 2012; Lammel *et al*., 2012; Yamaguchi *et al*., 2013; Ables, Park, and Ibañez-Tallon, 2023), which contribute to their functional segregation in regulating various behavioral processes (Kim and Chang, 2005; Namboodiri, Rodriguez-Romaguera, and Stuber, 2016; Trusel *et al*., 2019). In Pavlovian conditioning, the LHb plays a key role in negative prediction-error signaling (Bromberg-Martin and Hikosaka, 2011), temporal stability, and the extinction of conditioned fear memories (Bromberg-Martin and Hikosaka, 2011; Wang *et al*., 2013; Song *et al*., 2017; Durieux *et al*., 2020). Less is known about the MHb (Viswanath et al., 2013), although a few studies demonstrate that the MHb-IPN pathway regulates the expression of fear responses (Soria-Gómez et al., 2015; McLaughlin, Dani, and De Biasi, 2017) and their extinction (Zhang *et al*., 2016).

The MHb projects preferentially to the IPN through a dense fiber bundle known as the fasciculus retroflexus (FR) (Herkenham and Nauta, 1979; Hikosaka, 2010). Studies demonstrate that cholinergic afferents from the ventral MHb (vMHb) to the IPN co-release glutamate and acetylcholine (Qin and Luo, 2009; Ren *et al*., 2011). Interestingly, although GABA generally inhibits transmitter release via Gi-coupled GABA_B_ receptors (GABA_B_Rs) (Sakaba and Neher, 2003; Ulrich and Bettler, 2007), activation of GABA_B_Rs on MHb terminals is shown to drastically increase presynaptic calcium influx and release of glutamate, acetylcholine, and neurokinin B onto target postsynaptic IPN neurons (Zhang *et al*., 2016; Koppensteiner *et al*., 2024). Accordingly, GABA_B_R-mediated signaling in the MHb-IPN circuitry facilitates the extinction of conditioned fear memories (Zhang *et al*., 2016).

Lateralization represents a central theme in understanding the structural asymmetry of the habenula and its functional role in regulating various behavioral processes, including fear responses (Concha and Wilson, 2001; Agetsuma *et al*., 2010; Dreosti *et al*., 2014; Duboué *et al*., 2017). The zebrafish dHb-IPN pathway has clear morphological and molecular left-right asymmetry (Hong *et al*., 2013; Dreosti *et al*., 2014; Chen *et al*., 2019). Nodal-related laterality mutants, for example, *frequent-situs-inversus (fsi)* (Barth *et al*., 2005) and southpaw-deficient zebrafish (Long et *al.*, 2003), demonstrate how early left-right patterning can affect both visceral and neural asymmetries. These zebrafish mutants showed concordant reversal of the internal organ and diencephalic asymmetries. However, much less is known about a potential left-right asymmetry in the mammalian MHb, the mammalian homolog of the zebrafish dHb. Although a limited number of studies have shown morphological asymmetry in the dorsal habenula of smaller vertebrates (Gugliemotti and Fiorino, 1998; Concha and Wilson, 2001), the side-selective functional roles of the mammalian MHb-IPN pathway, based on confined manipulations of the left or right MHb, remain unexplored. Furthermore, no studies have compared the properties and regulation of synaptic transmission between the left and right pathways. In the present study, we investigated the synaptic properties of the left and right MHb inputs to the IPN in mice and addressed whether potential synaptic laterality in the MHb-IPN pathway results in its asymmetrical role in conditioned fear expression.

## Results

### Repetitive side-specific stimulation of MHb projections reveals bilateral input to single rIPN neurons

To investigate the synaptic features of the left and right MHb inputs to the IPN in adult mice, we employed 1-mm thick slices cut at a 53° angle to preserve the fasciculus retroflexus fiber bundle (FR) from the MHb to the IPN (Bhandari *et al*., 2021; Koppensteiner *et al*., 2024). This preparation allows selective electrical stimulation of the left and right MHb-derived axons and the recording of bilateral responses in a single postsynaptic IPN neuron (Fig. 1a). To rule out potential cross-stimulation, we stimulated the FR in one hemisphere with a 100-Hz train. This potentiated excitatory postsynaptic current (EPSC) responses only on the side receiving the 100-Hz stimulus, without influencing EPSCs evoked from the contralateral axons (**Fig. 1b**), thus confirming the stimulation’s confinement to the respective sides.

**Figure 1.**
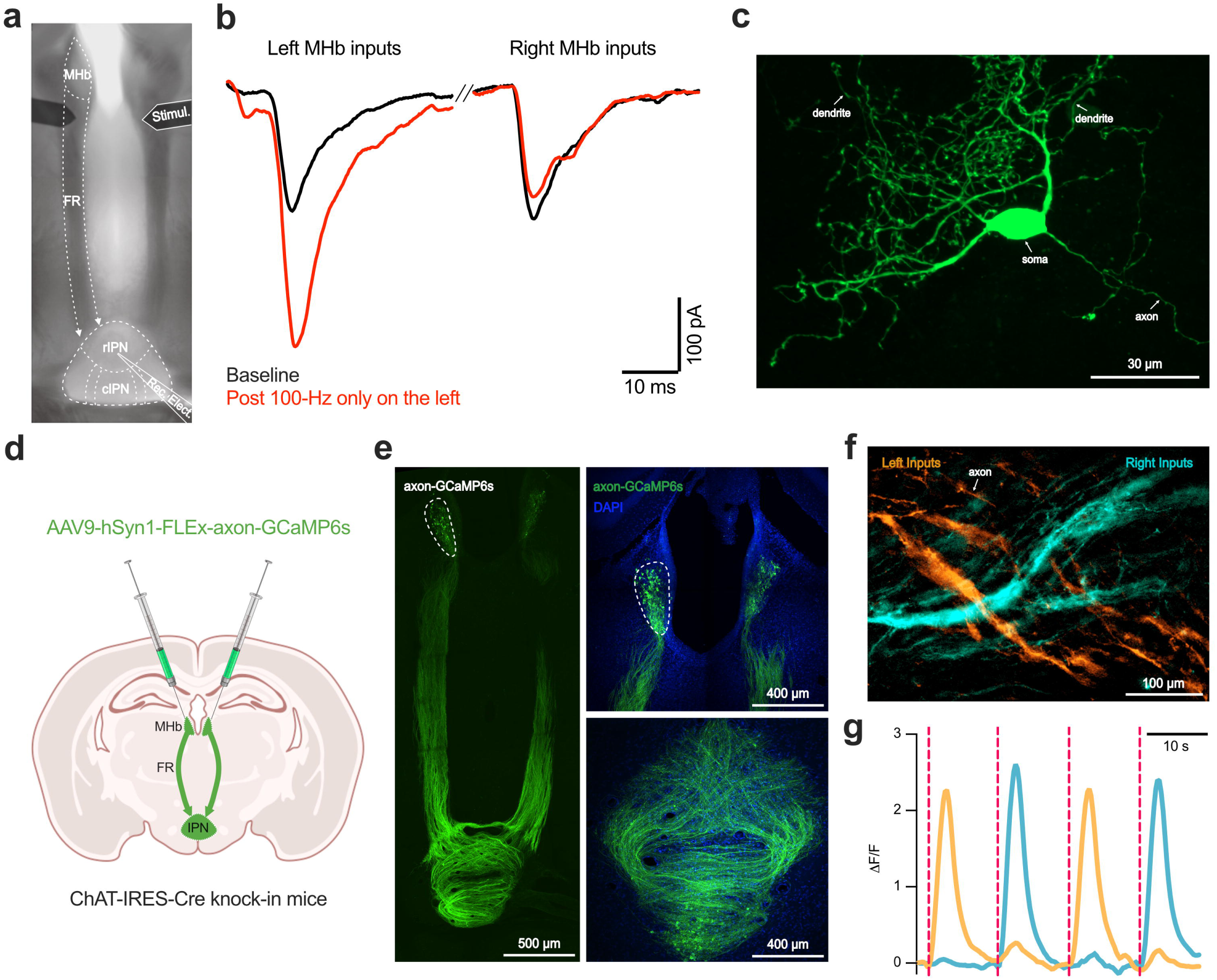
Selective recording from left and right MHb inputs in single IPN neurons. **a)** Representative image showing the experimental setup for the selective stimulation of left and right medial habenula (MHb) inputs at the fasciculus retroflexus (FR) and recording excitatory postsynaptic currents (EPSCs) in the same postsynaptic neuron in the rostral interpeduncular nucleus (rIPN) using 1 mm-thick slices. Concentric bipolar electrodes (Stimul.) are placed at the beginning of the FR, and a recording electrode is placed in the rIPN. **b)** First, the basal EPSCs were measured from left and right inputs individually. Next, only the left inputs were stimulated with 100-Hz high-frequency stimulation to induce potentiation at those synapses. Lastly, post-stimulation EPSCs were measured from left and right inputs again. EPSCs from the left but not right inputs were potentiated, indicating that the electrical stimulation of the left and right inputs was independent without crosstalk to the contralateral side. **c)** High magnification confocal image of a biocytin-filled IPN neuron showing the soma, axon, and intense dendritic arborization. **d)** Schematic illustration of the experimental design for AAV-mediated calcium imaging using GCaMP6s. AAV9-hSyn1-FLEx-axon-GCaMP6s was bilaterally injected into the MHb of ChAT-IRES-Cre knock-in mice to selectively label cholinergic neurons, enabling visualization of FR and comparison of Ca^2+^ responses in left and right MHb-derived axons. **e)** Representative confocal images of axon-GCaMP6s fluorescence in MHb cell bodies (top right), FR (left), and crossing axons in the IPN (bottom right). **f)** Rolling ball background subtracted fluorescent image showing crossing left and right MHb axons in the rIPN. Fibers were pseudo-colored for left and right inputs based on the standard deviation of the signal in response to 10-Hz trains. The image of the left MHb axon is the same data as published in Figure 2N (Koppensteiner et al., 2024). **g)** Fluorescence traces showing Δ*F/F* in response to 10-Hz electrical stimulation of the left and right FR based on responsive regions in **f,** indicating distinct, side-specific, and non-overlapping presynaptic calcium responses.

Interestingly, we noted that all the recorded neurons in the rostral subnucleus of the IPN (rIPN) received bilateral MHb inputs. This observation suggests that although the left and right MHb input fibers follow distinct and separate paths towards the IPN (**Fig. 1e**), MHb in both sides innervates all neurons within the rIPN. Intense dendritic arborization of IPN neurons allows them to receive bilateral inputs from both habenulae (**Fig. 1c**).

We next employed presynaptic calcium imaging further to rule out cross-communication between the left and right inputs. We injected viral vectors encoding an axon-enriched calcium indicator (AAV9-hSyn-DIO-axon-GCaMP6s-EGFP) into the bilateral ventral MHb of adult ChAT-IRES-Cre knock-in mice (**Fig. 1d**). Three weeks following the injections, GCaMP fluorescence revealed rich axons from both left and right MHb projecting to the rIPN in acute slice preparations (**Fig. 1e**). These axons traversed the midline in the rIPN, and extended across other IPN subnuclei, reaching the ventral region of the central nuclei while forming additional crossings (**Fig. 1e**). We then conducted calcium imaging in the rIPN where left and right MHb input fibers intersect while stimulating the incoming fiber paths using 10-Hz electrical stimulation. This approach provided a unique opportunity to observe both input fibers within the same field of view. We compared the presynaptic fluorescence intensity at rest and during stimulation as a metric of evoked activity in the region of interest (ROI) for each of the left and right stimuli (**Fig. 1f**). We found that stimulation of each side induced a robust increase in Ca^2+^ influx without significant overlap between the left and right inputs (**Fig. 1g**). The marginal response observed during the contralateral stimulation likely resulted from fibers running beneath the recorded surface. Overall, these data demonstrate that the left and right MHb input fiber stimulation had minimal crosstalk, thus providing further support for the convergence of bilateral MHb synaptic inputs in the IPN.

### Asymmetrical release probability in the MHb-derived inputs to the rostral IPN

To address potential side-dependent synaptic differences in glutamatergic projection originating from the vMHb to the rIPN, we first focused on synaptic release probability (P_r_), a crucial determinant of synaptic strength and plasticity (Schulz, Cook, and Johnston, 1994; Dobrunz and Stevens, 1997). We performed whole-cell recordings of rIPN neurons and stimulated left and right MHb fibers with paired pulses with a 50-millisecond inter-stimulus interval, ensuring a 6-second interval between opposite-side stimuli and a 10-second interval between consecutive same-side stimuli (**Fig. 2a**). Paired-pulse ratio (PPR), calculated as the ratio of the second evoked excitatory postsynaptic current (EPSC) to the first EPSC, serves as an index of synaptic P_r_, with higher PPR values generally reflecting lower P_r_ (Dobrunz and Stevens, 1997). Our recordings revealed significantly higher PPR in synapses derived from left vMHb terminals compared to those from the right side (*W = −689.0, n = 53 cells, p = 0.0019, Wilcoxon matched-pairs signed-rank test*; **Fig. 2b**), indicating a lower P_r_ in left vMHb-derived synapses. Notably, during recordings, we also observed that left vMHb synapses typically required several stimulations to achieve a stable release state, with a higher rate of failed synaptic release events (*W = −75, n = 17 cells, p = 0.0166, Wilcoxon matched-pairs signed-rank test*; **Fig. 2c**).

**Figure 2.**
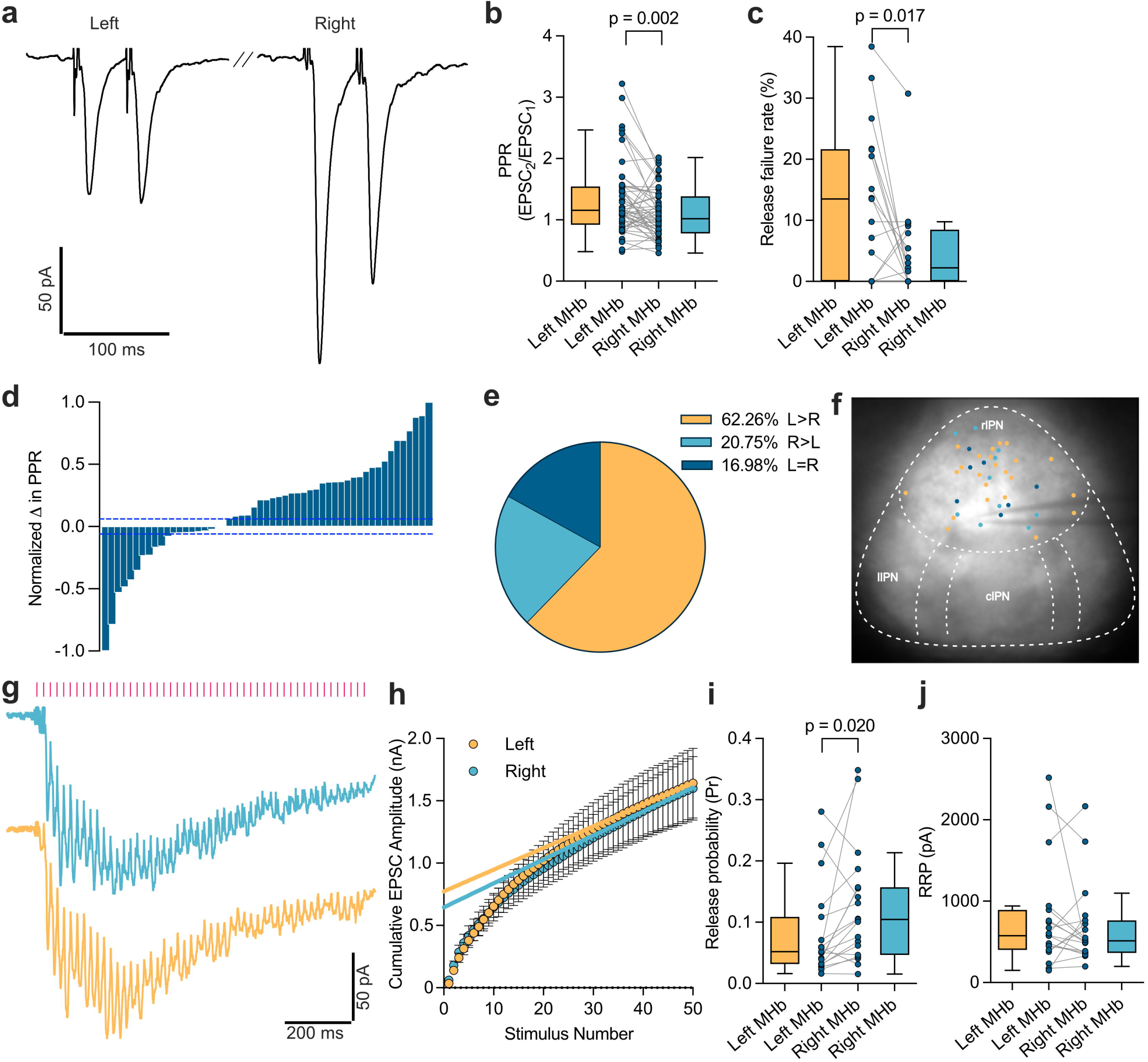
Asymmetrical release probability in MHb terminals. **a)** Example paired-pulse traces (50 ms interval) of EPSCs in the rIPN, evoked by electrical stimulation of the left and right FR. **b)** The paired-pulse ratio (PPR), calculated as EPSC2/EPSC1, showed that synapses derived from the left MHb exhibit a significantly higher PPR than those from the right MHb in the same postsynaptic rIPN neurons, indicating a lower release probability (P_r_) in left than right MHb terminals (*W = −689.0, n = 53 cells, p = 0.0019, Wilcoxon matched-pairs signed-rank test*). **c)** Synaptic release failure threshold was calculated as EPSCs lower than 4 times the pre-stimulus baseline current’s standard deviation. Paired comparison showing significantly higher rate of failed synaptic release events in the left compared to right MHb inputs in response to electrical stimulation (*W = −75.0, n = 17 cells, p = 0.0166, Wilcoxon matched-pairs signed-rank test*). **d)** Normalized differences in PPR between left and right inputs for each cell are plotted in a sequential order from the highest PPR difference for right MHb inputs (−1.0) to the highest PPR difference for left MHb inputs (1.0). The difference values are categorized using 15% of standard deviation (blue dashed line), within which are defined as equal PPRs. **e)** Pie chart showing the categorized PPRs. The majority of neurons in the rIPN show higher PPRs for left MHb inputs. **f)** Location of recorded neurons with their respective PPR category indicated by the same color as **e,** showing a homogenous distribution. **g)** Example of averaged EPSC traces for inputs originating from the left (orange) and right (blue) MHb following a 50-Hz stimulation (pink bars), demonstrating depletion of the readily releasable vesicle pool. **h)** Cumulative mean EPSC amplitudes plotted as a function of stimulus number for left and right MHb inputs (*n = 19 cells*). The error bars are presented as ± SEM. **i)** Paired comparison showing a significantly lower P_r_ for left than right MHb-derived synapses based on the first EPSC amplitude divided by the readily releasable pool size (RRP) using the high-frequency depletion protocol (*W = 114.0, n = 19 cells, p = 0.020, Wilcoxon matched-pairs signed-rank test*). **j)** Paired comparison of RRPs showing no significant difference between left and right MHb-derived synapses (*W = −20.0, n = 19 cells, p = 0.7086, Wilcoxon matched-pairs signed-rank test*). Box-and-whisker plots show the median, interquartile range (IQR), and whiskers extending to 1.5 times the IQR (Tukey method).

To determine whether the synaptic asymmetry was driven by the input side rather than the target cell location, we first categorized the recorded rIPN neurons based on the difference in RRPs. By normalizing the differences between the paired PPR values obtained with the left and right input stimulation and categorizing recorded neurons into three groups, we found that the majority of the rIPN cells have higher PPRs for the left input (**Fig. 2d-e**) (PPR L > R: 62.26%, L= R: 16.98%, L< R: 20.75%). To examine the location of neurons based on their PPR differences, we mapped the recorded neurons in rIPN with this categorization (**Fig. 2f**). Interestingly, the observed differences in P_r_ were independent of their location in the rIPN. The homogeneous distribution of neurons indicates that synaptic asymmetry arises from the input side rather than the target side.

To further verify the lower P_r_ in left MHb-derived synapses, we used a high-frequency (50 Hz) stimulation protocol to exhaust the readily releasable pool (RRP) of vesicles, thereby allowing us to analyze cumulative EPSC amplitudes over successive stimuli (**Fig. 2g-h**). (Schneggenburger, Meyer and Neher, 1999). We confirmed that left vMHb-derived synapses have significantly lower P_r_ relative to right vMHb-derived synapses in the rIPN (*W = 114.0, n = 19 cells, p = 0.020, Wilcoxon matched-pairs signed-rank test*; **Fig. 2i**), while no significant difference was detected in the RRP size (**Fig. 2j**).

### Calcium-permeable AMPARs are present at similar levels in synapses receiving left and right MHb inputs

Next, to test whether synaptic asymmetry also involves postsynaptic mechanisms, we compared calcium-permeable AMPA receptors (CP-AMPARs) in synapses receiving left- or right-MHb inputs. These receptors have previously been reported to contribute to the long-lasting enhancement of glutamate release in the MHb-IPN pathway (Koppensteiner, Melani, and Ninan, 2017). We recorded EPSCs at holding potentials of −60, −40, −20, 0, +20, +40, and +60 mV using an internal solution containing spermine (**Fig. 3a**), and plotted the current/voltage relationship (**Fig. 3b**). The slopes from −60 to 0 mV, and those from 0 to +60 mV were calculated to obtain rectification indexes for the left and right MHb-derived synapses. Inwardly rectifying currents, characteristic of CP-AMPARs, were observed in both of them (**Fig. 3c**, *main effect of slopes: F_(1, 22)_ = 25.10 p< 0.0001, main effect of side: F_(1, 22)_ = 0.5104 p = 0.4825, side and slope interaction: F_(1, 22)_ = 0.05207 p=0.8216*), with Sidak post-hoc correction (*p = 0.0054 and p = 0.0025 for left and right MHb slopes, respectively*) with comparable levels of rectification indexes (**Fig. 3d**). These rectification indexes are consistent with previous findings (Koppensteiner, Melani and Ninan, 2017), and demonstrate that the thick slice-cutting method followed by long-range stimulation accurately captures the synaptic characteristics of the MHb-IPN pathway.

**Figure 3.**
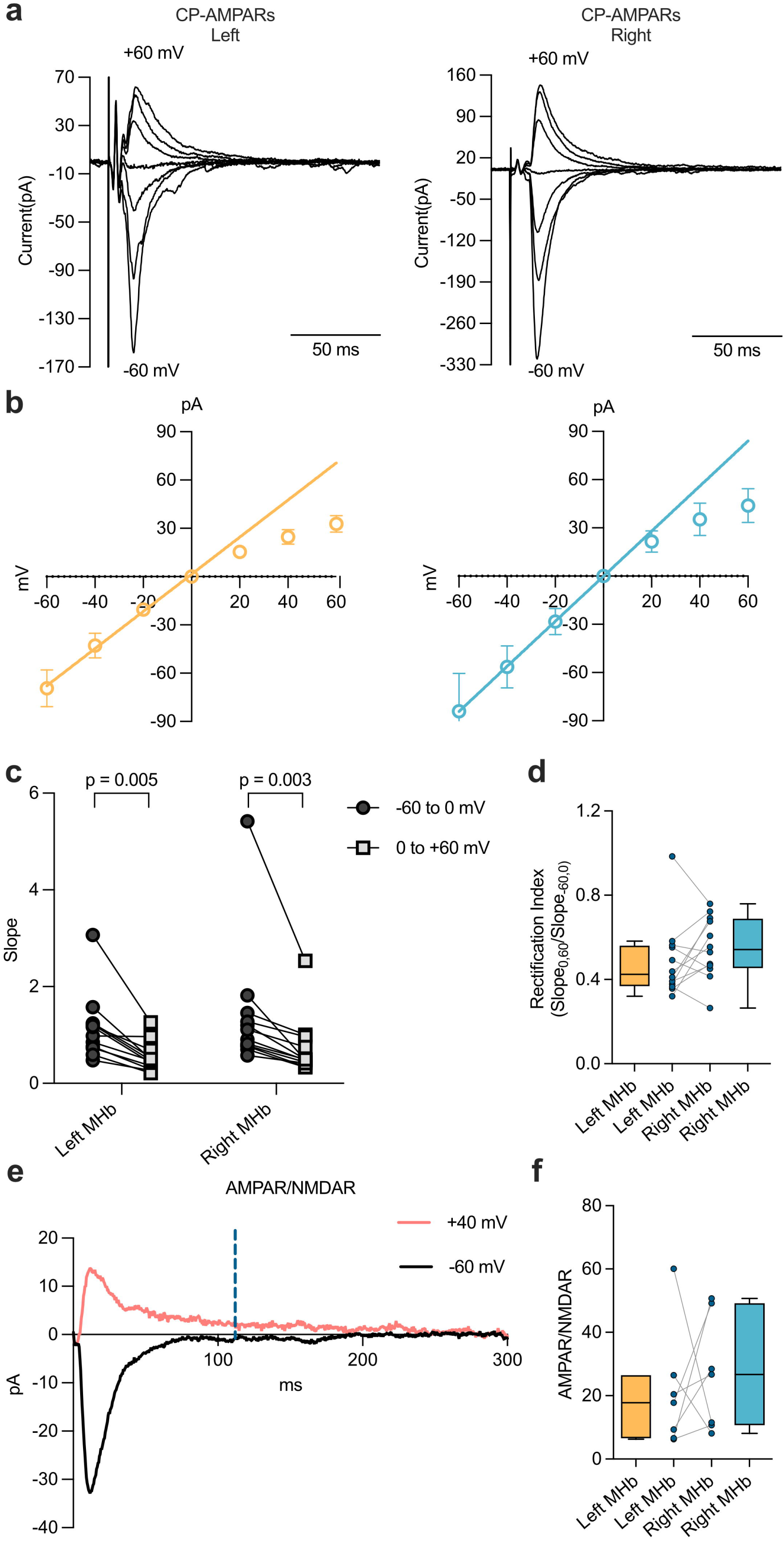
No difference in postsynaptic properties in left and right MHb projections. **a)** Representative current-voltage (I-V) traces recorded from neurons receiving left and right MHb-derived inputs. The traces show EPSCs evoked at different holding potentials (−60 to +60 mV) in the presence of spermine in the intracellular solution. **b)** I-V plots showing the characteristic inwardly rectifying current, indicative of calcium-permeable AMPA receptors (CP-AMPARs) for both left (orange) and right (blue) MHb-derived synapses. **c)** Comparison of slopes from −60 mV to 0 mV and those from 0 mV to +60 mV calculated for left and right MHb-derived inputs. The data indicate significant inward rectification in synapses from both sides, confirming the presence of CP-AMPARs. Two-way ANOVA showed a main effect of negative/positive slopes, (*main effect of slopes: F_(1, 22)_= 25.10 p< 0.0001, main effect of side: F_(1, 22)_= 0.5104 p=0.4825, side and slope interaction: F_(1, 22)_= 0.05207 p=0.8216*), with Sidak post-hoc correction (*p = 0.0054 and 0.0025 for left and right MHb slopes, respectively*). **d)** Connected pairs showing no difference in the rectification index between neurons receiving left and right MHb inputs (*W = 34.0, n = 12 cells, p = 0.2036, Wilcoxon matched-pairs signed-rank test*). **e)** Representative traces showing AMPAR current at −60 mV (black) and NMDA receptor (NMDAR) current at +40 mV holding potentials. The vertical, blue-dashed line indicates the time point for measuring the isolated NMDAR component. **f)** Graph depicting the AMPA/NMDA ratio in neurons receiving left and right MHb inputs. The data suggest NMDARs are present on both sides, though at low levels with no difference between left and right MHb-derived synapses (*W = 8.0, n = 7 cells, p = 0.5781, Wilcoxon matched-pairs signed-rank test*). Box-and-whisker plots show the median, interquartile range (IQR), and whiskers extending to 1.5 times the IQR (Tukey method).

We also examined the AMPA/NMDA ratios by recording EPSCs at −60 mV and +40 mV holding potentials to assess potential side-dependent asymmetry. The NMDA currents were measured at 100 milliseconds after the AMPA peak current (**Fig. 3e**). We observed marginal NMDAR-mediated components with no difference in the AMPA/NMDA ratios between left and right MHb-derived inputs (**Fig. 3f**). Based on the comparable levels of CP-AMPARs and AMPA/NMDA ratios between left and right inputs recorded in the same postsynaptic neurons, we concluded that the input-side dependent asymmetry in synaptic transmission is ascribable to presynaptic mechanisms.

### Presynaptic GABA_B_R activation differentially potentiates left and right MHb-derived inputs

Previous studies have demonstrated that baclofen, a GABA_B_R agonist, potentiates synaptic transmission in the MHb-IPN pathway by increasing Ca²LJ influx at presynaptic terminals (Zhang *et al*., 2016; Koppensteiner, Melani, and Ninan, 2017; Koppensteiner *et al*., 2024). To explore whether this potentiation shows input-side dependent difference, we stimulated left and right-derived fiber paths individually and recorded EPSC amplitudes in the absence and presence of baclofen (1 µM) (**Fig. 4a**). The relative EPSC amplitudes, obtained by dividing the EPSC amplitudes after baclofen application with the basal EPSC amplitudes, showed significantly higher degree of GABA_B_R-mediated potentiation for the left than right inputs (**Fig. 4b**, *W = −272.0, n = 27 cells, p = 0.0006, Wilcoxon matched-pairs signed-rank test*) despite having similar basal release amplitudes (**Fig. 4c**). The PPRs in the presence of baclofen showed no difference between left and right MHb inputs, indicating an equalization of P_r_ (**Fig. 4d**). One notable observation was the complete abolishment of failures to evoke EPSC responses during GABA_B_R-mediated potentiation in contrast to the basal state (**Fig. 4e**, *Left MHb: W = −78, n = 17 cells, p = 0.0005; Right MHb: W = −55, n = 17 cells, p = 0.0020, Wilcoxon matched-pairs signed-rank test*). Altogether, these data indicate that the left MHb-IPN pathway may play a key role in plasticity mechanisms for emotion regulation, while the right MHb pathway predominates in the basal state.

**Figure 4.**
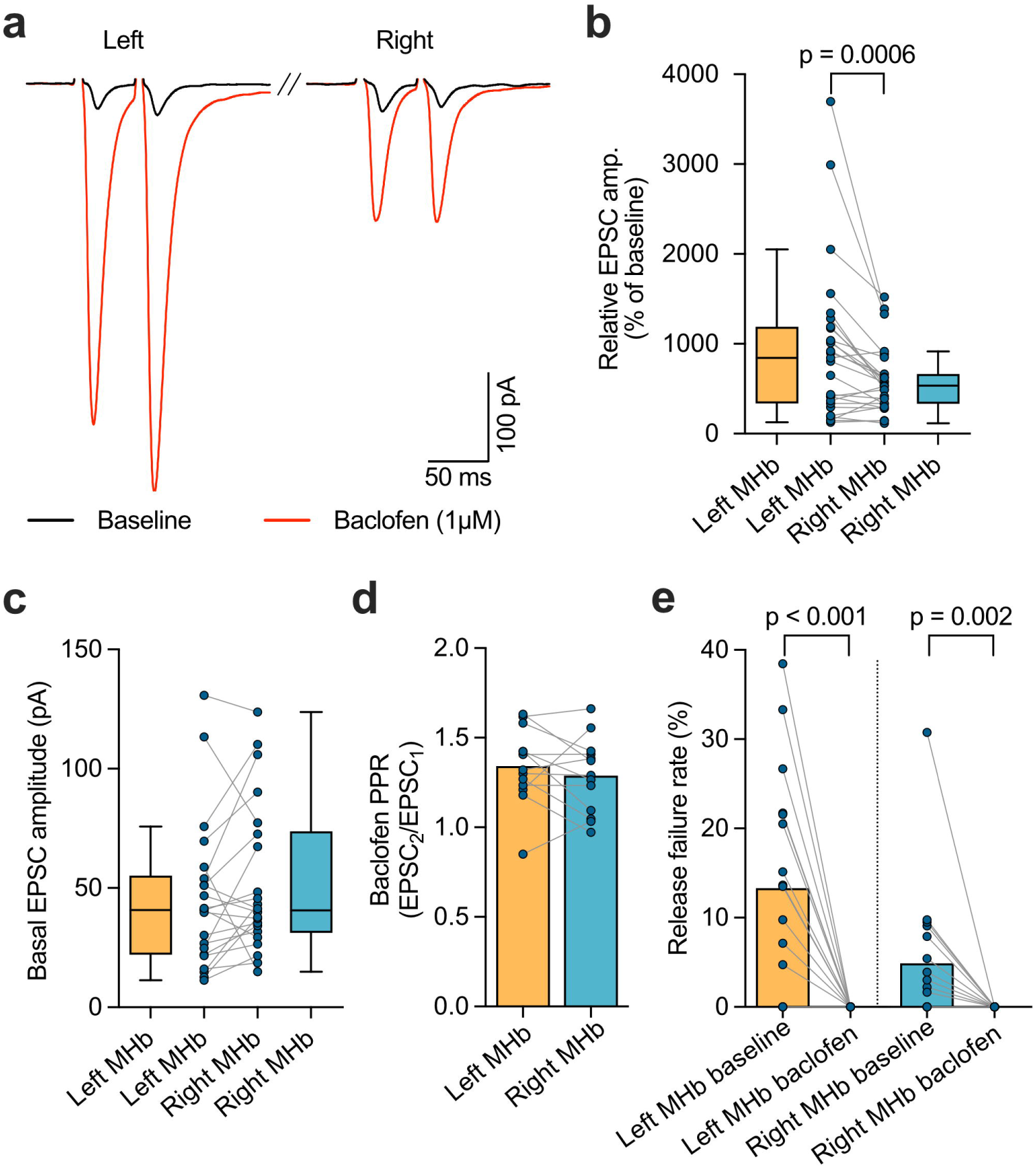
Asymmetrical GABA_B_R-mediated potentiation in left and right MHb synapses. **a)** Representative traces showing EPSCs recorded for left and right MHb-derived inputs before (black) and after (red) the application of baclofen (1 µM), a GABA_B_ receptor (GABA_B_R) agonist. The traces indicate drastic potentiation in response to GABA_B_R activation, with left inputs showing a greater relative EPSC increase compared to right inputs on the same postsynaptic neuron. **b)** Box plot with connected pairs showing the relative increase in EPSC amplitudes following baclofen treatment in left and right MHb-derived synapses in the rIPN. A significantly higher relative potentiation was found in left than right MHb-derived synapses (*W = −272.0, n = 27 cells, p = 0.0006, Wilcoxon matched-pairs signed-rank test*). **c)** No difference in the basal EPSC amplitudes for the left and right MHb-derived synapses in the same postsynaptic IPN neurons (*W = 95, n = 22 cells, p = 0.1289, Wilcoxon matched-pairs signed-rank test*). **d)** Comparison of the PPR between left and right MHb-derived inputs following baclofen application, showing similar PPRs following GABA_B_R activation (*t = 1.018, df = 12, n = 13 cells, p = 0.3288, paired t-test*). **e)** Bar graph showing complete abolishment of failed synaptic release events during baclofen treatment. The baseline data are the same as shown in Fig. 2b. The decrease in failure rate was more prominent in the left inputs (*Left MHb: W = −78, n = 17 cells, p = 0.0005; Right MHb: W = −55, n = 17 cells, p = 0.0020, Wilcoxon matched-pairs signed-rank test*). Box-and-whisker plots show the median, interquartile range (IQR), and whiskers extending to 1.5 times the IQR (Tukey method).

### Lateralization persists in i.v. situs solitus but not in i.v. situs inversus

In zebrafish, the nodal signaling dictates the structural left-right asymmetry in the internal organs as well as in the dHb-IPN pathway (Long et *al.*, 2003; Barth *et al*., 2005). To determine whether embryonic left-right patterning by the nodal signaling underlies the functional asymmetry in MHb-to-IPN synapses in mice, we examined PPR and baclofen sensitivity from inversus viscerum (*i.v.*) mutants exhibiting either situs solitus or situs inversus. In *i.v.* animals with situs solitus, the PPR measurement confirmed that left MHb terminals maintain a lower P_r_ than the right, similar to the wild-type (WT) mice (*W = −186.0, n = 24 cells, p = 0.0065, Wilcoxon matched-pairs signed-rank test*; **Fig. 5a, left**). However, in *i.v.* animals displaying situs inversus, there was no significant difference in PPR between the left and right inputs (*W = −102.0, n = 27 cells, p = 0.2268, Wilcoxon matched-pairs signed-rank test*; Fig. **5a****, right**).

**Figure 5.**
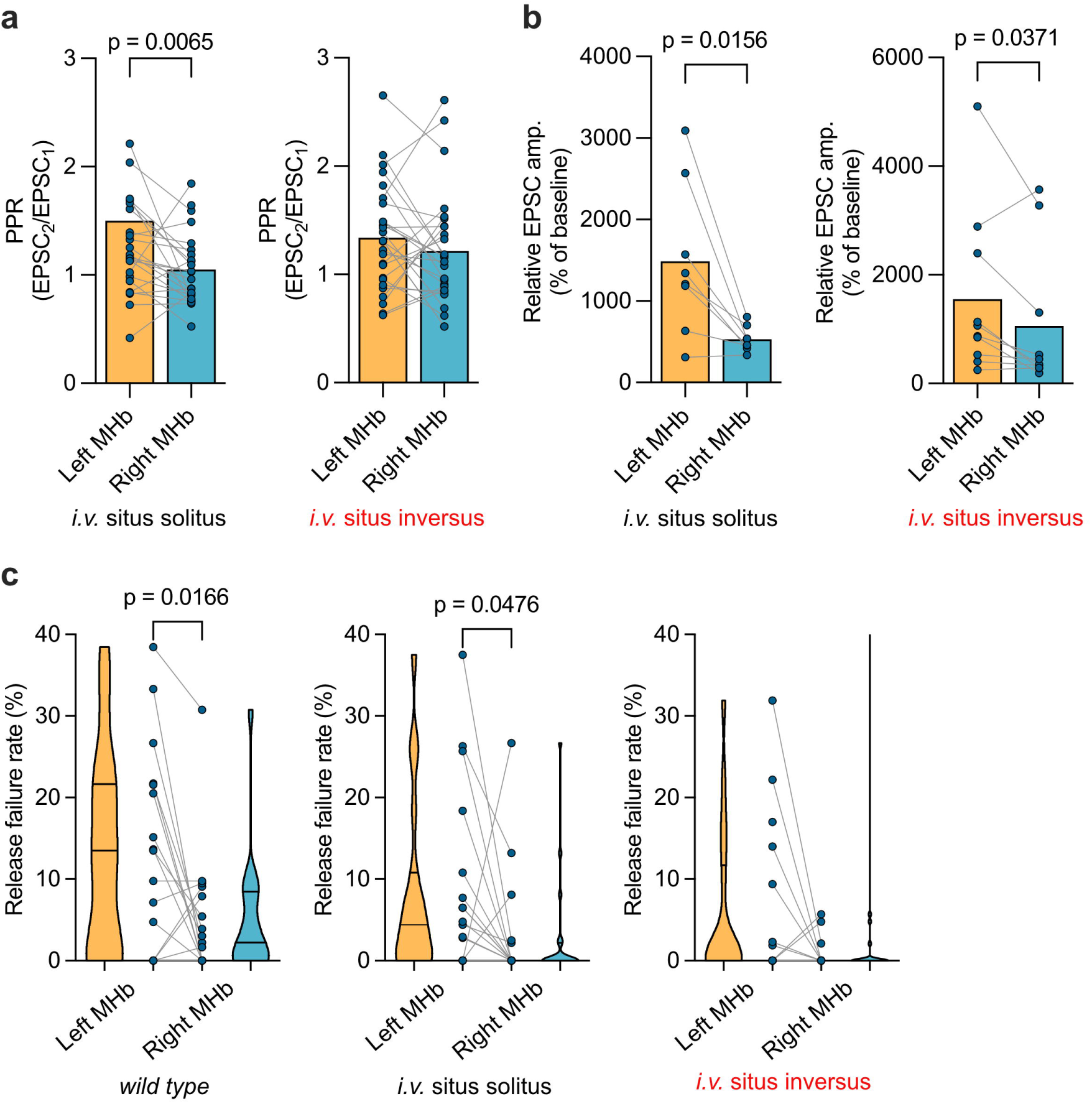
Release probability and release failure rate comparison between situs solitus and situs inversus *i.v.* mutant mice. **a**) The PPR indicates a lower release probability (P_r_) in left than right MHb inputs in *i.v.* situs solitus mutants similar to WT mice (*W = −186.0, n = 24 cells, p = 0.0065, Wilcoxon matched-pairs signed-rank test*), but no difference in *i.v.* situs inversus mutants (*W = −102.0, n = 27 cells, p = 0.2268, Wilcoxon matched-pairs signed-rank test*). **b)** A significantly higher GABA_B_R-mediated relative EPSC amplitude increase was found in left than right MHb-derived synapses in *i.v.* situs solitus mutants, similar to that in WT mice (*W = −34.0, n = 8 cells, p = 0.0156, Wilcoxon matched-pairs signed-rank test).* A marginal significance in the same direction as situs solitus was detected in situs inversus mutants (*W = −41.0, n = 10 cells, p = 0.0371, Wilcoxon matched-pairs signed-rank test*). **c)** Violin plots with connected pairs showing differences in release failure rates for left and right MHb-derived inputs. Significant difference between left and right inputs in WT mice (same as Fig. 2c) is shown for comparison. A lesser, but significant difference was detected in situs solitus mutants (*W = −63.0, n = 19 cells, p = 0.0476, Wilcoxon matched-pairs signed-rank test*), whereas no difference was found in situs inversus mutants (*W = −13.0, n = 17 cells, p = 0.4961, Wilcoxon matched-pairs signed-rank test)*. Violin plots show the minimum and maximum values and the quartiles.

We next examined the GABA_B_R-mediated potentiation in these mutants. As in WT mice, situs solitus animals showed a significantly higher potentiation ratio on the left side than on the right (*W = −34, n = 8 cells, p = 0.0156, paired t-test*; **Fig. 5b, left**). In contrast, *i.v.* mutants with situs inversus displayed a marginal significance for left-right difference (*W = −41.0, n = 10 cells, p = 0.0371, Wilcoxon matched-pairs signed-rank test*; **Fig. 5b, right**).

Finally, we compared the failure rates of the release event between left and right MHb-derived synapses in *i.v.* mutant mice. Higher failure rate was found in left than right MHb-derived synapses in situs solitus mice (*W = −63.0, n = 19 cells, p = 0.0476, Wilcoxon matched-pairs signed-rank test*; **Fig. 5c, middle**), though the significance was marginal compared to that in WT mice (WT, *W = −75.0, n = 17 cells, p = 0.0166, Wilcoxon matched-pairs signed-rank test*; **Fig. 5c, left,** the same data as **Fig. 2c**). However, no difference in failure rate between input-sides was found in situs inversus mice (*W = −13.0, n = 17 cells, p = 0.4961, Wilcoxon matched-pairs signed-rank test*; **Fig. 5c, right**). Since we found no sign of the reversed asymmetry in situs inversus animals, these results suggest that the defect of the nodal signaling distinctly affects the asymmetry in the internal organs and the MHb-IPN pathway in mice, in contrast with its effect in zebrafish.

### Chemogenetic inhibition of the left but not the right vMHb cholinergic neurons attenuates the recall of cued fear memories

Based on our findings showing a significantly larger GABA_B_R effect on left than right MHb-derived inputs, we hypothesized that synaptic asymmetry may be involved in shaping behavioral functions. We focused on conditioned fear memories, based on previous evidence showing a critical role for the MHb in controlling fear responses (Agetsuma *et al*., 2010; Zhang *et al*., 2016; Duboué *et al*., 2017). To this end, we employed ChAT-IRES-Cre knock-in mice that were unilaterally injected with the Cre-dependent inhibitory designer receptor exclusively activated by designer drugs (iDREADDs) virus into the left or right MHb (**Fig. 6a and b; Suppl. Fig. 1a**). This approach allowed us to achieve targeted silencing of cholinergic neurons via activation of iDREADDs with clozapine-N-oxide (CNO). We first verified the efficacy of the inhibition by recording neuronal firing activity in acute brain slices derived from ChAT-IRES-Cre knock-in mice expressing the virus (**Suppl. Fig. 1b**). Whole-cell voltage clamp and cell-attached recordings of mCherry-positive vMHb neurons showed that CNO administration significantly suppressed neural firing compared with the basal state (**Suppl. Fig. 1c**), and its washout recovered the firing to the baseline state (**Suppl. Fig. 1c and 1d**).

**Figure 6.**
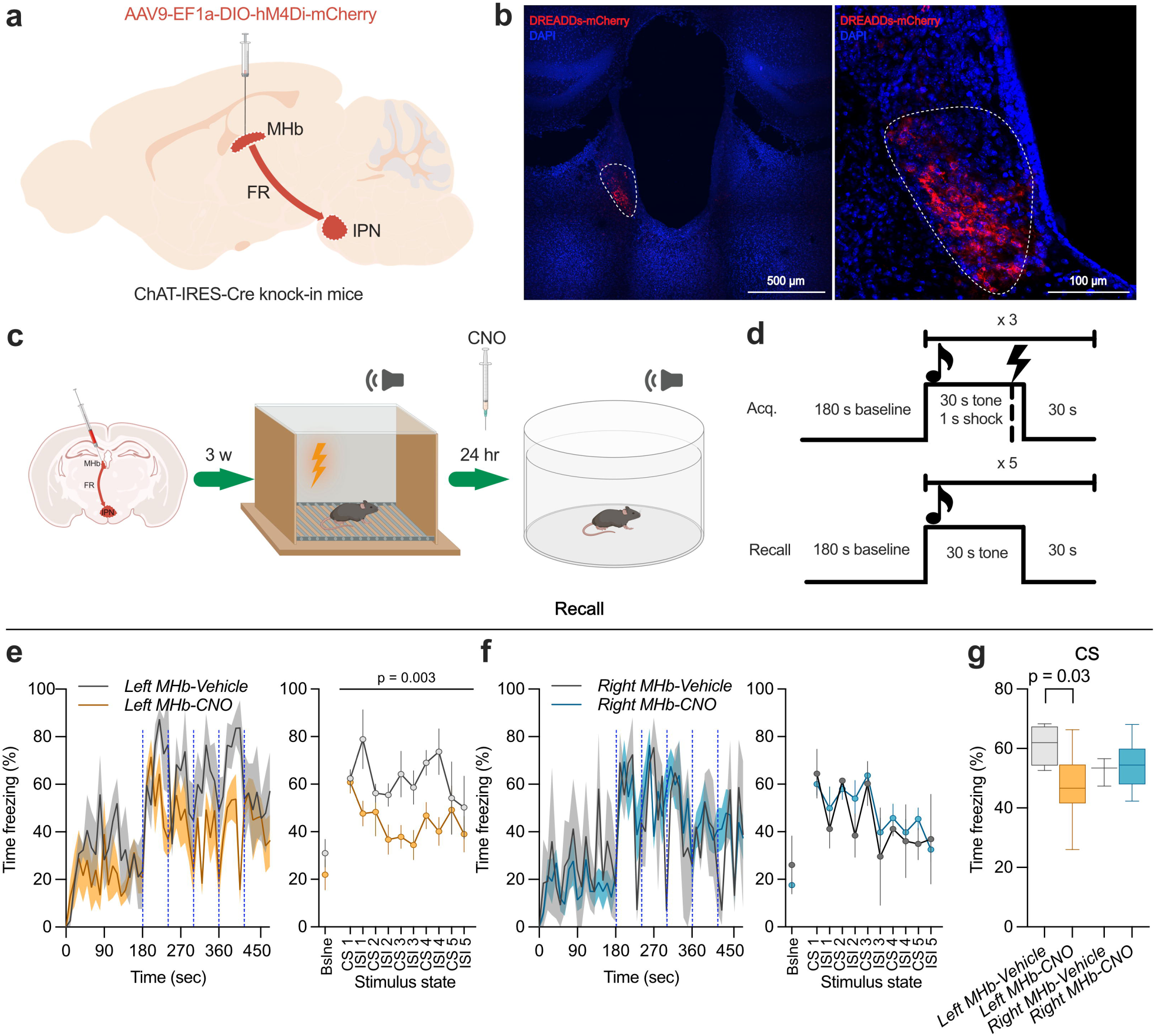
Chemogenetic inhibition of the left but not the right MHb cholinergic neurons attenuates fear response. **a)** Schematic representation of the experimental design, showing the targeted chemogenetic inhibition of cholinergic neurons in the left or right MHb using AAV9-EF1a-DIO-hM4Di-mCherry virus in ChAT-IRES-Cre knock-in mice. **b)** Representative confocal images of the injection site of the AAV in the vMHb, showing strong DREADD-mCherry expression (red) in the cholinergic vMHb neurons. **c)** Diagram showing the timeline of the viral injection, cued fear conditioning in context A on day 1, and retrieval in context B on day 2. CNO or vehicle (saline) was applied 24 hours after the conditioning and an hour before the recall, followed by re-exposure to the tone to assess freezing behavior. **d)** The fear conditioning protocol. Acquisition (Acq.) involved 3 minutes of baseline followed by five co-terminating tone-shock pairings. Recall was performed 24 hours later, with mice exposed to the tone without shock. **e, f)** Time courses of freezing behavior in 10-second bins during recall following chemogenetic inhibition of the left MHb (**e**, left panel, orange) or the right MHb (**f**, left panel, blue), compared to vehicle controls. Vertical blue dashed lines indicate the presentation of the conditioned stimuli (CS, tone). Inhibition of the left but not the right MHb reduced freezing time compared with the control. All groups show low freezing levels in the baseline habituation phase in context B. Over the CS and inter-stimulus intervals (ISI), left MHb inhibition (orange) shows significantly reduced freezing compared to the control (**e**, right panel), whereas right MHb inhibition (blue) does not significantly alter freezing behavior (**f**, right panel) (*main effect of side: F_(1,18)_ = 1.008, p= 0.329; main effect treatment: F_(1,18)_ = 3.559, p= 0.075; side and treatment interaction: F_(1,18)_ = 8.059 p= 0.011; post-hoc comparison, Left MHb-vehicle vs Left MHb-CNO: p = 0.003; Right MHb-vehicle vs Right MHb-CNO: p = 0.525, repeated measures ANOVA with Tukey’s post-hoc comparison*). The error bars are presented as ± SEM. **g)** Boxplot showing the freezing time percentage during the CS presentation. Inhibition of the left but not right MHb significantly reduced fear expression compared to vehicle controls (*main effect of side: F_(1,18)_ = 0.038, p = 0.848, main effect of treatment: F_(1,18)_ = 1.827, p = 0.193, side and treatment interaction: F_(1,18)_ = 3.404, p = 0.082, post-hoc comparison, Left MHb-vehicle vs Left MHb-CNO, p = 0.03; Right MHb-vehicle vs Right MHb CNO: p = 0.741, Two-Way ANOVA with Tukey’s post-hoc comparison*). Left MHb-vehicle, n = 4; Left MHb-CNO, n = 7; Right MHb-vehicle, n = 3; Right MHb-CNO, n = 8. The error bars indicate ± SEM. Box-and-whisker plots show the median, interquartile range (IQR), and whiskers extending to the minimum and maximum values. See also Supplementary Figures S1 and S2.

Next, we employed the cued fear-conditioning paradigm, in which adult ChAT-IRES-Cre knock-in mice expressing hM4Di-mCherry in the left or right vMHb were conditioned to associate a mild aversive foot shock with a discrete tone (**Fig. 6c and 6d**). For all groups, freezing responses increased across tone-shock trials (*main effect of stimulus: F_(3.038, 54.683)_ = 31.857 p < 0.001, repeated measures ANOVA (Greenhouse-Geisser corrected*), (*Left MHb-vehicle, n = 4; Left MHb-CNO, n = 7; Right MHb-vehicle, n = 3; Right MHb-CNO, n = 8,* ***Suppl.* Fig. 2a and 2b**). One day after the conditioning, CNO (i.p. 3 mg/kg) or vehicle was injected into the mice 40 min before testing for cued memory recall in a different neutral context (Context B). Animals from all experimental groups showed minimal baseline freezing levels, indicating that the new context itself was not aversive and that the chemogenetic manipulation did not induce the generalization of fear responses. Freezing behavior increased upon presentation of the conditioned discrete tone stimulus in all groups (**Fig. 6e-g**). However, the magnitude of fear responses was significantly weaker in the group expressing hM4Di in the left MHb compared with the vehicle control group (**Fig. 6e**, right, time course of freezing levels to the CS and during inter-stimulus intervals (ISI), *main effect of side: F_(1,18)_ = 1.008 p= 0.329, main effect treatment: F_(1,18)_ = 3.559 p= 0.075, side and treatment interaction: F_(1,18)_ = 8.059 p= 0.011; post-hoc comparison, Left MHb-vehicle vs Left MHb-CNO: p = 0.003, repeated measures ANOVA with Tukey’s post-hoc comparison*, **Fig. 6g**, time freezing during tone-CS, *main effect of side: F_(1,18)_ = 0.038, p = 0.848, main effect treatment: F_(1,18)_ = 1.827 p = 0.193, side and treatment interaction: F_(1,18)_ = 3.404 p = 0.082; post-hoc comparison, Left MHb-vehicle vs Left MHb-CNO, p = 0.03, Two-Way ANOVA with Tukey’s post-hoc comparison*), indicating that the acquired fear memory was not effectively expressed upon inhibition of left MHb neurons. In contrast, CS-evoked freezing levels in the group expressing hM4Di in the right MHb did not differ from those observed in the respective vehicle control group (**Fig. 6f, right,** time course of freezing levels to the CS and during ISI, *Right MHb-vehicle vs Right MHb-CNO: p = 0.525,* **Fig. 6g**, time freezing during tone-CS, *Right MHb-vehicle vs Right MHb CNO: p = 0.741*). Taken together, the findings indicate that the vMHb is functionally lateralized in mice, and that the left MHb cholinergic neurons may exert a facilitatory role on cued fear memory recall.

### Knockout of GABA_B_Rs in cholinergic neurons of the left but not the right MHb attenuates fear memory recall

Finally, given that left MHb inputs show lower P_r_ but higher GABA_B_R-mediated potentiation (**Fig. 4**), we investigated whether GABA_B_R signaling in the mouse MHb-IPN circuitry is involved in the lateralized modulation of fear responses. We utilized Cre-expressing lentiviruses (LV-hSyn1-Cre-mCherry) to achieve conditional knockout of GABA_B_Rs in the left or right MHb of adult GABA_B_1 floxed mice (**Fig. 7a**). Three weeks following the virus delivery, we conducted immunohistochemical evaluation on brain slices derived from injected mice to verify the efficacy of the procedure. In the control group of GABA_B_1 floxed mice injected with mCherry-expressing virus (LV-hSyn1-mCherry), targeted neurons co-expressed mCherry and GABA_B_R (**Fig. 7b, left**). By contrast, in the Cre-mCherry group, the complete absence of GABA_B_R expression in mCherry-positive cell bodies confirmed successful knockout of GABA_B_Rs in the targeted neurons (**Fig. 7b, right**). Visualization of MHb axonal terminals in the IPN showed that while mCherry-expressing axons in Cre-mCherry-injected mice were engulfed by GABA_B_R1-expressing fibers, no co-localization of the two signals was observed (**Fig. 7c**), further confirming the selective knockout of GABA_B_Rs in the targeted presynaptic terminals.

**Figure 7.**
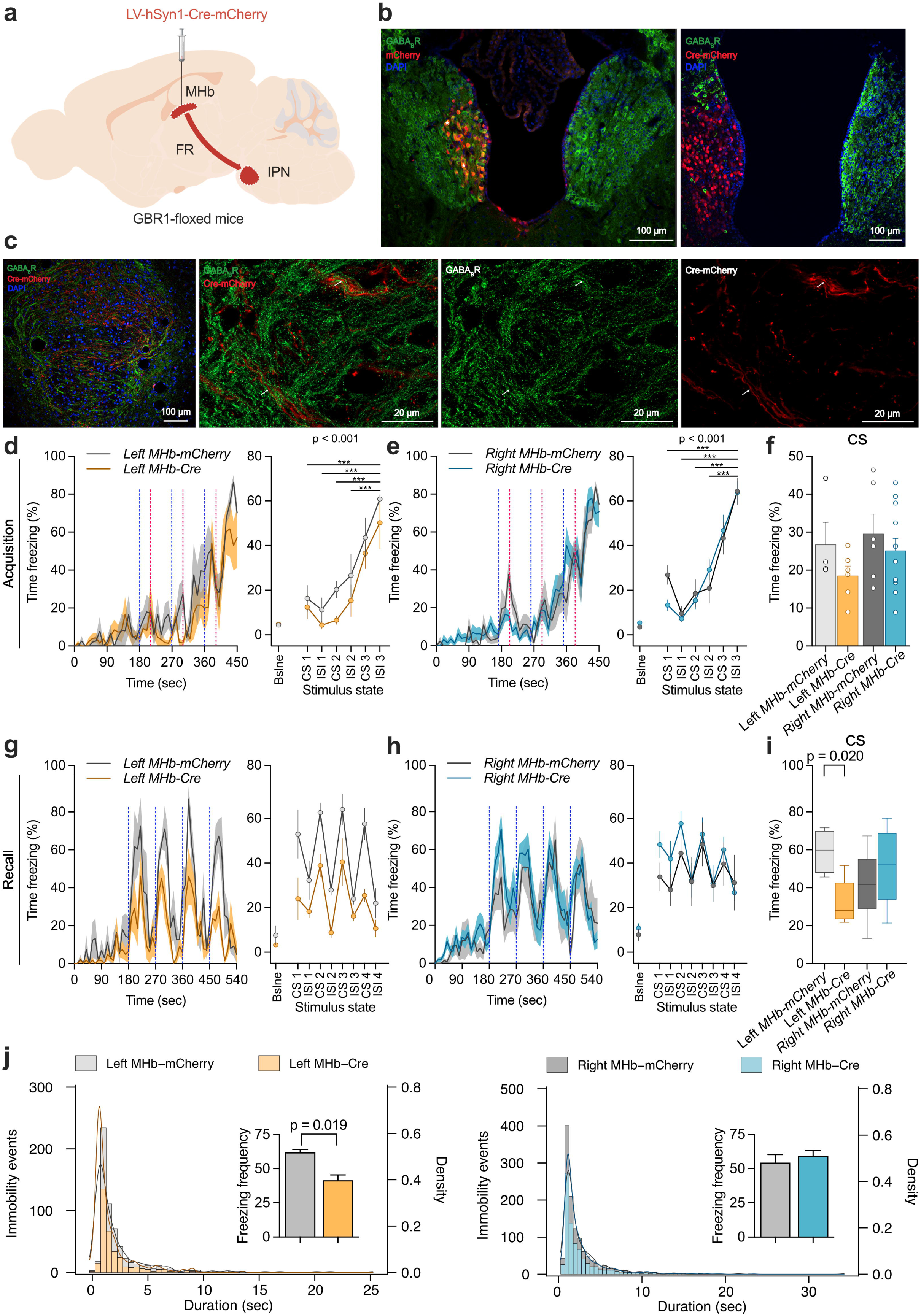
Conditional knock-out of GABA_B_1 in left MHb neurons attenuates cued fear response. **a)** Schematic of the experimental design for conditional GABA_B_ receptor (GABA_B_1) knock-out in left or right MHb neurons using the Cre-lox system and lentivirus-mediated (LV) Cre expression. Mice were injected with either a control virus (LV-hSyn1-mCherry) or a Cre-expressing virus (LV-hSyn1-Cre-mCherry) to conditionally knock out GABA_B_1 unilaterally in the targeted MHb neurons. The diagram illustrates the MHb projection to the IPN via the FR. **b)** Representative immunohistochemistry images showing successful knockout of GABA_B_1 in the targeted ventral MHb neurons. In the control virus-infected MHb neurons, mCherry (red) is co-expressed with GABA_B_1 (green) (left), whereas in Cre-expressing virus-infected neurons, no GABA_B_1 immunoreactivity is observed in mCherry-positive cell bodies (right). **c)** Images of MHb axons and terminals in the rIPN showing mCherry and GABA_B_1 labeling (a low magnification image on the left). Higher magnification images demonstrate the absence of GABA_B_1 immunoreactivity (green) in mCherry-Cre-expressing axons (red). **d, e)** Time courses of freezing behavior in 10-second bins during the fear acquisition phase in GABA_B_1 floxed (GBR1-floxed) mice unilaterally injected with Cre-expressing lentivirus (LV-hSyn1-Cre-mCherry) or mCherry-expressing lentivirus (LV-hSyn1-mCherry) as a control. Both left MHb-Cre (**d**, orange, left panel) and right MHb-Cre (**e**, blue, right panel) groups show increasing freezing time over CS and ISI, indicating successful acquisition of fear memory (*main effect of stimulus: F_(3.318, 79.622)_ = 56.174, p < 0.001, repeated measures ANOVA (Greenhouse-Geisser corrected)*). **f)** The percentages of freezing time during the conditioned stimulus (CS) presentation in the fear acquisition phase exhibit no significant difference between groups (*main effect of side: F_(1, 24)_ = 1.192, p = 0.286; main effect of virus: F_(1, 24)_ = 2.098, p = 0.160, side and virus interaction: F_(1, 24)_ = 0.188, p = 0.668; post-hoc comparison, Left MHb-Cre vs Left MHb-mCherry: p = 0.245; Right MHb-Cre vs Right MHb-mCherry: p = 0.416; repeated measures ANOVA with Tukey’s post-hoc comparison*). The error bars indicate ± SEM. **g, h)** Time courses of freezing behavior in 10-second bins during the cued fear recall phase. Vertical blue dashed lines indicate the CS presentation. All groups show low levels of freezing in the baseline habituation phase in context B. Over the CS and ISI, conditional GABA_B_1 knock-out mice in the left MHb (orange) tended to reduce freezing behavior compared to the control mice, whereas those in the right MHb (blue) does not alter freezing behavior compared to the control (*main effect of side: F_(1, 24)_ = 0.849, p = 0.366, main effect of virus: F_(1, 24)_ = 1.090, p = 0.307, side and virus interaction: F_(1, 24)_ = 3.927, p = 0.059; post-hoc comparison, Left MHb-Cre vs Left MHb-mCherry: p = 0.068*, *Right MHb-Cre vs Right MHb-mCherry: p = 0.452 repeated measures ANOVA with Tukey’s post-hoc comparison*). **i)** Boxplot showing the freezing time percentage during CS. Conditional GABA_B_1 knock-out in the left but not right MHb significantly reduced freezing behavior compared to the control (*main effect of side: F_(1, 24)_ = 0.009, p = 0.926; main effect of virus: F_(1, 24)_ = 1.601, p = 0.218, side and virus interaction: F_(1, 24)_ = 7.180, p = 0.013; post-hoc comparison, Left MHb-Cre vs Left MHb-mCherry: p = 0.020; Right MHb-Cre vs Right MHb-mCherry: p = 0.260; Two-Way ANOVA with Tukey’s post-hoc comparison*). **j)** All immobility event durations plotted as a histogram showing the number of immobility events per bin and the density curve (left panel, left MHb; right panel, right MHb). Density plot showing distinct freezing durations around 0.9 seconds, which is used as a freezing threshold. Bar graph showing the left MHb-Cre group decreased the frequency of freezing episodes to CS and ISI rather than shortening individual episodes. (*main effect of side: F_(1, 24)_ = 0.988, p = 0.330; main effect of virus: F_(1, 24)_ = 2.297, p = 0.143, side and virus interaction: F_(1, 24)_ = 6.055, p = 0.021; post-hoc comparison, Left MHb-Cre vs Left MHb-mCherry: p = 0.019; Right MHb-Cre vs Right MHb-mCherry: p = 0.448; repeated measures ANOVA with Tukey’s post-hoc comparison)*. Left MHb-mCherry, n = 4; Left MHb-Cre, n = 6; Right MHb-mCherry, n = 6; Right MHb-Cre n = 12. Box-and-whisker plots show the median, interquartile range (IQR), and whiskers extending to the minimum and maximum values. See also Supplementary Figure S3

To address the relevance of GABA_B_Rs in the MHb for fear memory, we employed the cued fear-conditioning paradigm (**Suppl. Fig. 3a-b**) again. During conditioned acquisition, all groups, regardless of the virus or injected side, showed a gradual increase in freezing responses with consecutive tone-shock pairings (**Fig. 7d, e**, right, *main effect of stimulus: F_(3.318, 79.622)_ = 56.174, p < 0.001, repeated measures ANOVA (Greenhouse-Geisser corrected), Left MHb-mCherry, n = 4; Left MHb-Cre, n = 6; Right MHb-mCherry, n = 6; Right MHb-Cre, n = 12*). Importantly, since targeted GABA_B_R knockout was conducted prior to testing, successful acquisition of the CS-US association indicated that targeted GABA_B_R knockout in the left or right MHb does not interfere with the ability to acquire fear memories. When tested for fear memory recall in a neutral context (Context B), mice that received conditional GABA_B_R knockout in the left MHb demonstrated significantly less freezing than those in the mCherry group following the presentation of the CS (**Fig. 7g**, right, time course of freezing levels to the CS and during ISI, *side effect: F_(1, 24)_ = 0.849, p = 0.366, virus effect: F_(1, 24)_ = 1.090, p = 0.307, side and virus interaction: F_(1, 24)_ = 3.927, p = 0.059; post-hoc comparison, Left MHb-Cre vs Left MHb-mCherry: p = 0.068*, *repeated measures ANOVA with Tukey’s post-hoc comparison;* **Fig. 7i**, time freezing during tone-CS, *main effect of side: F_(1, 24)_ = 0.009, p = 0.926, main effect of virus: F_(1, 24)_ = 1.601, p = 0.218, side and virus interaction: F_(1, 24)_ = 7.180, p = 0.013; post-hoc comparison, Left MHb-Cre vs Left MHb-mCherry: p = 0.020, Two-Way ANOVA with Tukey’s post-hoc comparison*). By contrast, conditional GABA_B_R knockout in the right MHb did not change cued fear recall compared with the mCherry control (**Fig. 7h**, right, time course of freezing levels to the CS and during ISI, *Right MHb-Cre vs Right MHb-mCherry: p = 0.452*, **Fig. 7i**, time freezing during tone-CS, *Right MHb-Cre vs Right MHb-mCherry: p = 0.260*). To gain a better understanding of the fear memory phenotype, we evaluated the distribution of immobility events throughout the recall session (**Fig. 7h**). We observed an increased number of immobility events around 0.9 milliseconds, which we used as a minimum freezing duration threshold. Additionally, we found that decreased fear responses in left MHb-Cre mice stemmed from a decrease in the number of freezing episodes rather than a shortening of the duration of individual freezing bouts (**Fig. 7j**, *side effect: F_(1, 24)_ = 0.988, p = 0.330, virus effect: F_(1, 24)_ = 2.297, p = 0.143, side and virus interaction: F_(1, 24)_ = 6.055, p = 0.021; post-hoc comparison, Left MHb-Cre vs Left MHb-mCherry: p = 0.019; Right MHb-Cre vs Right MHb-mCherry: p = 0.448, repeated measures ANOVA with Tukey’s post-hoc comparison*). These outcomes support the results following chemogenetic inhibition of the left and right cholinergic MHb (**Fig. 6e-f**). Overall, our findings highlight lateralization of the mouse MHb, with the left MHb playing a key role in modulating conditioned fear responses via GABA_B_R signaling.

## Discussion

Our study reveals a previously unknown lateralization in the MHb-IPN pathway in the mouse. While individual rIPN neurons receive bilateral inputs from the left and right MHb, these inputs differ in fundamental ways. The left MHb synapses exhibit lower P_r_ under basal conditions and display a more pronounced potentiation upon GABA_B_R activation than the right MHb synapses, ultimately influencing fear memory expression. Those asymmetrical synaptic properties remain in situs solitus but not situs inversus *i.v.* mutant mice, showing dissociation of asymmetry between the internal organs and the MHb-IPN pathway. Chemogenetic inhibition or knockout of GABA_B_1 receptors, selectively in the left but not the right MHb, impairs conditioned fear responses, establishing a direct link between these asymmetric synaptic properties and lateralized behavioral modulation.

### Synaptic Asymmetry in Release Probability

Our findings reveal a lower basal P_r_ in the left MHb-IPN pathway compared to the right side (**Fig. 2)**, a feature typically associated with enhanced synaptic facilitation and greater temporal summation during high-frequency activity (Atwood and Karunanithi, 2002). The observed asymmetry in P_r_ likely arises from intrinsic presynaptic properties, independent of the postsynaptic side. Molecular differences in proteins involved in vesicle docking and priming in the active zone, such as RIM and Munc13, may also play a role (Augustin *et al*., 1999; Kaeser *et al*., 2011). For instance, reduced RIM-mediated coupling of calcium channels to vesicle release sites or less efficient Munc13-dependent vesicle priming in the left MHb could result in lower basal P_r_ while allowing greater dynamic modulation during synaptic activity. Another reason for this asymmetry in P_r_ may arise from differences in synaptotagmin (Syt) isoforms, which regulate calcium-dependent vesicle release. Syt-1, an ubiquitously expressed isoform, is a fast calcium sensor driving synchronous, high-probability vesicle release, enabling rapid responses during brief, high-frequency signaling (Fernández-Chacón *et al*., 2001). In contrast, Syt-9, which is strongly and specifically expressed in the habenula (Allen Brain Atlas), favors asynchronous or slower release, particularly contributing to the enhanced short-term facilitation during repetitive stimulation (Xu, Mashimo, and Südhof, 2007). In the left MHb terminals, a higher ratio of Syt-9 to Syt-1 expression than in the right MHb terminals could underlie their lower P_r_ and greater facilitation potential during sustained activity. A striking property of the MHb-IPN pathway is the prominent presynaptic localization of the R-type voltage-gated Ca^2+^ channel (VGCC) subunit Cav2.3 (Parajuli et al., 2012), which solely triggers vesicular release in MHb terminals (Zhang et al., 2016; Bhandari et al., 2021). Differential localization of Cav2.3 in the presynaptic active zone of left and right MHb terminals could also underlie the asymmetry in P_r_.

### Asymmetry in GABA_B_R-mediated Potentiation

GABA_B_Rs regulate neurotransmitter release primarily by inhibiting VGCCs, activating G-protein-coupled inwardly rectifying potassium (GIRK) channels, reducing vesicle priming, or regulating the fusion machinery at many synapse types (Chalifoux and Carter, 2011; Gassmann and Bettler, 2012). In contrast to the common inhibitory role of GABA_B_Rs, previous studies have shown that GABA_B_R activation in the MHb-IPN pathway can paradoxically increase neurotransmitter release (Zhang *et al*., 2016; Koppensteiner *et al*., 2024). Our findings extend this peculiar property by demonstrating that the magnitude of GABA_B_R-mediated potentiation is greater for left than right MHb inputs (**Fig. 4**). GABA_B_Rs are abundantly expressed in the peri-synaptic region of MHb terminals with lower densities inside the active zone, and co-localized with Cav2.3 in virtually all MHb terminals (Bhandari et al., 2021). Differences in GABA_B_R density and the distance to Cav2.3 between the left and right MHb terminals could affect the efficacy of this modulation. Otherwise, a larger space for the potentiation due to the lower P_r_ in the left than right MHb terminals may result in a higher effect of GABA_B_R activation even with the same GABA_B_R localization and coupling with Cav2.3. Although the source of GABA for physiological activation of GABA_B_R is still unclear, most of the IPN neurons are GABAergic (Antolin-Fontes et al., 2015), making it likely that strong excitatory MHb inputs increase the ambient GABA concentration, activating GABA_B_R in MHb terminals (Koppensteiner et al., 2017; Zaupa et al., 2021). This situation could enable greater dynamic modulation of the left MHb inputs during high activity of GABAergic IPN neurons. GABA_B_R-mediated facilitation may also involve molecular adaptations that differ between hemispheres. Auxiliary proteins such as KCTD subunits are known to modulate GABA_B_R kinetics and sensitivity (Rajalu *et al*., 2015). KCTD proteins accelerate GABA_B_R desensitization, fine-tuning their ability to modulate synaptic transmission. A differential expression or subunit composition of these proteins in the left versus right MHb could contribute to the observed asymmetry in GABA_B_R-mediated potentiation. Additionally, we showed that GABA_B_R activation can recruit additional vesicles into the RRP (Koppensteiner *et al*., 2024). In the left MHb, recruitment of more vesicles with higher P_r_ might disproportionately enhance synaptic efficacy, effectively equalizing P_r_ between hemispheres during GABA_B_R activation. The two-pool hypothesis of vesicle dynamics (Koppensteiner *et al*., 2024; Guzikowski and Kavalali, 2024) further supports this model, assuming that distinct vesicle subpopulations within the RRP are mobilized during basal versus GABA_B_R-mediated transmission. Physiologically, a stronger GABA_B_R-mediated potentiation in the left MHb may serve as a modulatory “gain control” mechanism. When the circuit demands heightened excitability, such as during the recall of a fear memory, this left-sided potentiation rapidly boosts synaptic output, facilitating the retrieval of aversive memories.

### Effect of *i.v.* Mutation on Habenular Asymmetry

We showed that *i.v.* mutants with situs solitus but not situs inversus exhibit similar left-right differences in P_r_ and, to a lesser degree, GABA_B_R-mediated potentiation as WT mice. This finding supports the idea that normal left-sided nodal signaling is crucial for establishing normal lateralization in the MHb, as observed in zebrafish. By contrast, situs inversus mutants failed to show the normal or reversed left-right difference in P_r_ and GABA_B_R-mediated potentiation, resulting in no asymmetry, suggesting that nodal signaling that reversed the internal organ laterality does not have a strong enough influence to reverse the laterality of the MHb-IPN pathway. This result implies that the default mode of synaptic properties is symmetrical, consistent with our previous findings in the CA3-CA1 hippocampal pathway in mice (Kawakami et al., 2008). This pathway also shows input-side dependent asymmetry in WT mice (Kawakami et al., 2003; Shinohara et al., 2008), but the asymmetry disappeared in the *i.v.* mutant mice. However, in this pathway, the loss of asymmetry occurred in both situs solitus and situs inversus animals (Kawakami et al., 2008), indicating a much weaker influence of nodal signaling than in the MHb-IPN pathway. Our results show that the failure rate of left MHb-derived synapses decreased in situs solitus mice compared to WT mice, and even further decreases in situs inversus mice, getting closer to that in right MHb-derived synapses in WT mice, indicating that nodal flow disruption reduces the left-sided differentiation in the MHb-IPN pathway in a similar way as we found in the right isomerism mice in the hippocampus (Kawakami et al., 2008). Nonetheless, the MHb in mice also shows similarities to the zebrafish dHb, with left-right patterning paralleling the visceral axis at least in situs solitus animals.

### Behavioral Implications of Lateralized Synaptic Function

Our behavioral experiments revealed that chemogenetic inhibition of the left MHb significantly attenuates cued fear memory recall, whereas inhibition of the right MHb had no such effect (**Fig. 6**). Similarly, conditional knockout of GABA_B_1 in the left MHb impaired fear memory recall, reinforcing the notion of a left-specific role of the MHb in fear expression (**Fig. 7**). This lateralization appears to be specific to the recall phase, as memory acquisition was not affected by these manipulations. These findings suggest that fear encoding and storage might be bilaterally represented, whereas recall involves lateralized processing in the left MHb. Bilateral manipulation studies of MHb have shown that enhancing cholinergic inputs reduces fear-conditioned freezing (Soria-Gómez et al., 2015) and that disrupting MHb inputs impairs fear extinction (Zhang *et al*., 2016) in mice, somewhat contrasting with our unilateral findings. This discrepancy underscores the importance of considering lateralization in experimental designs. The lateralization observed in the MHb may serve to optimize neural processing by assigning specific functions to each hemisphere, thereby enhancing the efficiency of fear memory recall. This specialization could facilitate rapid and precise responses to aversive stimuli, which are critical for survival. The specialization of left MHb for recall might reflect an evolutionary adaptation to minimize redundancy and enhance processing efficiency. Studies in avian and amphibian habenulae demonstrate that lateralized circuits reduce overlap while facilitating parallel processing of distinct behavioral functions (Rogers, 2002). Such division of labor could be particularly advantageous in fear circuits, where precise and rapid recall of aversive stimuli is crucial for survival (Güntürkün, Ströckens, and Ocklenburg, 2020).

Input-side-dependent asymmetries have also been documented in other brain regions, notably the hippocampus (Kawakami et al., 2003; Shinohara et al., 2008; Kohl *et al*., 2011; Shipton et al., 2014). Specifically, synapses in the CA1 area made by the left CA3 pyramidal cells have denser NR2B subunit of the NMDA receptor, less GluA1 subunit of the AMPA receptor, smaller PSD size, and greater long-term potentiation than those made by the right CA3 cells. This hemispheric specialization within the hippocampus has been implicated in the left CA3-specific role in long-term spatial memory formation (Shipton et al., 2014). Interestingly, these findings parallel ours in the MHb, where left-sided inputs exhibit enhanced synaptic potentiation and impact on the behavior. These input-side-dependent asymmetries across different brain regions suggest that presynaptic origin can dictate synaptic properties on the same postsynaptic neuron, underscoring common principles and mechanisms underlying asymmetry formation.

Although lateralization of the habenula has been well established in non-mammalian species such as zebrafish, definitive evidence in mammals remained elusive. Our findings thus provide an evolutionary bridge, suggesting that lateralized habenular circuits may be a conserved principle across vertebrates. Furthermore, this study demonstrates, for the first time, a functional synaptic asymmetry in the MHb-IPN pathway. Our findings have raised important questions for the future. First, which molecular and cellular mechanisms underlie the asymmetry in P_r_ and GABA_B_R-mediated potentiation? Second, which specific downstream pathways are affected by left MHb activity, and does the found asymmetry extend to other emotional or cognitive behaviors? Third, is the input-side-dependent presynaptic and postsynaptic lateralization an extensive feature of the brain, influencing synaptic functions and behavior? Understanding these asymmetries provides valuable insights into the neural basis of the well-known lateralization of higher brain functions and may inform therapeutic strategies for neurological disorders characterized by dysregulated asymmetry.

## Materials and Methods

### Animals and ethics

Experiments in this study were performed using 8- to 16-week-old adult mice. The wild-type mice used were C57BL/6J (#000664) and ChAT-IRES-Cre knock-in (#006410) from Jackson Laboratory. The *i.v.* mutant mice had the SI/Col/C57BL/6J background as reported in Goto et al., 2010. For creating GABA_B_R conditional knockout mice via Cre-expressing lentivirus (LV), GABA_B_1^lox511/lox511^ floxed mice (Haller *et al*., 2004) were kindly provided by Bernhard Bettler (University of Basel, Switzerland). Mice were housed together and single-caged after the operation in a room with a 12-hour light/dark cycle and had ad libitum access to food and water. Replacement, reduction, and refinement (3R) principles were applied whenever possible. All experiments were carried out strictly following European animal experimentation guidelines and approved by the Bundesministerium für Wissenschaft, Forschung und Wirtschaft of Austria under the license number “BMWFW-66.018/0005-WF/V/3b/2015”.

### Stereotaxic virus injection and viral vectors

Stereotaxic injection of viruses and tracers was performed as previously described (Cetin *et al*., 2006). Mice were anesthetized by intraperitoneal injection of ketamine (100 mg/kg)/xylazine (4.5 mg/kg) and placed in a stereotaxic frame (Kopf). Metacam (5 mg/kg) and novalgin (33 mg/kg) were injected subcutaneously. The animal’s head was shaved, and lidocaine 3% was applied locally. The parietal skin of the head was cut with scissors to expose the skull; the periosteum was then removed, and the skull surface was cleaned. A nanoliter injector pump (World Precision Instruments, Sarasota, FL, USA) with a glass pipette was tilted 20° laterally and aligned on top of the bregma as a zero point for the injection coordinates. The injector was then moved to the coordinates of the region of interest respective to the brain atlas (Paxinos and Franklin, 2019). For MHb, −1.45 A/P, 0.60 L, −2.65 D/V, and for IPN −3.05 A/P, 0.1 L, −4.7 D/V were used as coordinates. A small burr hole was drilled over the region of interest, and the dura mater was exposed. The glass pipette was moved to the dura, and the brain surface was used as the zero point for dorsoventral coordinates. 200-500 nL of virus solution was loaded into the pipette, then the pipette was lowered to the injection site. The total volume was injected with a 50 nL/min flow rate, and the pipette was left in place for an additional 10 minutes. After infusion, it was slowly removed, and the skin was closed using surgical glue. For all the adeno-associated vectors, we used the AAV9 serotype. pAAV-hSynapsin1-FLEx-axon-GCaMP6s was a gift from Lin Tian (Addgene plasmid # 112010; http://n2t.net/addgene:112010; RRID:Addgene_112010) (Broussard *et al*., 2018), pAAV-hSyn-DIO-hM4D(Gi)-mCherry was a gift from Bryan Roth (Addgene plasmid # 44362; http://n2t.net/addgene:44362; RRID:Addgene_44362) (Krashes *et al*., 2011), and pAAV-hSyn-DIO-mCherry were from Bryan Roth (Addgene plasmid # 50459; http://n2t.net/addgene:50459; RRID:Addgene_50459). pLV-SYN1-Cre-P2A-mCherry and pLV-SYN1-P2A-mCherry were constructed by VectorBuilder and used for LV production in-house.

### Acute slice electrophysiology

Mice were anesthetized by intraperitoneal injection of an overdose ketamine/xylazine and transcardially perfused with ice-cold oxygenated (95% o_2_. 5% Co_2_) artificial cerebrospinal fluid (ACSF) containing (in mM): 118 NaCl, 2.5 KCl, 1.25 NaH_2_PO_4_, 1.5 MgSO_4_, 1 CaCl_2_, 10 Glucose, 3 Myo-inositol, 30 Sucrose, 30 NaHCO_3_; pH = 7.4 as described previously (Bhandari *et al*., 2021; Koppensteiner *et al*., 2024). The brain was carefully removed and placed in the cutting chamber of a vibratome (DSK7, Dosaka). A 1-mm slice containing the IPN, the FR, and the two habenulae was cut at 56° and placed into a recovery chamber containing standard ACSF at 37 LJ for 1 hr. Subsequently, the slice was moved to the recording chamber and continuously superfused with 2.5 mM CaCl_2_-containing, oxygenated ACSF at room temperature (RT), containing 20 µM bicuculline methiodide (Tocris, Bristol, UK), 5 µM mecamylamine hydrochloride, and 50 µM hexamethonium bromide. Stimulating electrodes were placed on top of the MHb cell bodies or the fiber tract, allowing the selective stimulation of either the left or right side. For a reliable comparison, stimulating electrodes were placed at the same level on the MHb-IPN axis in all experiments. To eliminate selection bias, we randomized the stimulating electrodes to the left and right sides, as well as the recording protocol. Patch pipettes (3-6 MΩ) were filled with the internal solution containing (in mM): 130 K-Gluconate, 10 KCl, 2 MgCl_2_, 2 MgATP, 0.2 NaGTP, 0.5 EGTA, 10 HEPES, 5 QX_314_-Cl; pH 7.4 adjusted with KOH. Electrophysiological signals were acquired at 10-50 kHz and filtered at 2.0 kHz using a Multiclamp 700B amplifier connected to a Digidata 1440A digitizer (Molecular Devices, CA, USA). Access resistance (Ra) was continuously monitored throughout all recordings, and cell changes in Ra exceeding 20% were discarded.

PPR experiments were performed using concentric bipolar electrodes for electrical stimulation. Electrodes were placed at the beginning of the FR at similar distances from the IPN, and the stimulation protocol and electrodes were shuffled between sides to avoid bias. A 0.2-millisecond pulse duration with a 50-millisecond inter-stimulus interval was used.

The protocol consisted of a 6-second interval between sides and a 10-second interval within the same side. Failure rates were calculated using the first pulse of the paired stimuli for both left and right inputs in the same trace. For each cell, the standard deviation of the baseline current (measured over a 2-s window between left and right stimulations) was computed across all traces; four times the found standard deviation was used as the threshold for defining a response as a “failure.” Any EPSC below this threshold was deemed a failed release event. The failure ratio was then computed as the number of failed events divided by the total number of traces, both under basal conditions and during baclofen treatment.

For 50-Hz high-frequency vesicular pool-depletion experiments, cumulative EPSC amplitudes were plotted, and a line was fitted to the last 10-15 stable responses. The RRP size was estimated by extrapolating the line to the first stimulus. P_r_ was estimated from the average EPSC amplitude and RRP size using the following equation: I p X q X N, where p is the P_r_, q is the quantal size, and N is the number of release sites. Finally, P_r_ was calculated as the average of the first EPSC amplitude divided by the RRP size.

### Acute slice calcium imaging

Acute slice calcium imaging was performed as previously described (Koppensteiner *et al*., 2024), and the data in this study were reused to analyze left and right MHb input identification in the rIPN. Three weeks following the AAV9-hSynapsin1-FLEx-axon-GCaMP6s bilateral injection into the MHb, thick acute brain slices were prepared as described previously. The slice was recovered at 37 ℃ for 40 min. We specifically centered the axon crossings such that the inputs from left and right MHb were visible in the same field of view. Using the same recording solution as for electrophysiological recordings, the slice was superfused with ACSF containing 2.5 mM Ca^2+^ at RT. Bilateral stimulating electrodes were placed at the beginning of the FR. Next, using a 20X 0.5 NA water-immersion objective (Olympus), MHb axons on the surface entering the rIPN were carefully focused, and the GCaMP expression was confirmed using blue LED excitation (pE-300 LED light source (CoolLED) and a monochrome CCD camera (XM10, Olympus). To visualize the fluorescent signal, excitation filter 460 – 490 nm, dichroic mirror 505 nm, and barrier filter 510 nm (U-MWIB2, Olympus) were used. GCaMP fluorescence in response to 10-Hz electrical stimulation (3-s duration, 0.2-ms pulse width, 0.5 – 2.5 V stimulation intensity) was recorded with a baseline period without any stimulation. Stimulation was alternated between the left and right sides at 3 Hz, and illumination was turned off between sweeps to prevent photobleaching. To minimize background noise, we employed a rolling-ball subtraction algorithm followed by standard deviation differentiation to identify fibers responsive to alternating stimulation from left and right inputs.

### Cued fear conditioning (CFC)

Mice were handled for 3 consecutive days before the experiment. For chemogenetic inhibition using ChAT-IRES-Cre knock-in mice, on the day of conditioning, animals were placed in a conditioning chamber (Context A) and, after three minutes, three conditioned stimuli (CS) (auditory tone 4 kHz, 80 dB, 30 s with 30 s interval) were presented with co-terminating unconditioned stimuli (foot shock 0.75 mA, 1 s with 1 m interval). At the end of the acquisition session, animals were removed from the conditioning chamber and returned to their home cages. Twenty-four hours later, mice were injected with CNO (3 mg/kg i.p.) or vehicle. 40 minutes after the injection, mice were placed in a different chamber (Context B) and received the CS without foot shock five times. Either left or right MHb cholinergic neurons were silenced using inhibitory DREADDs before the recall phase. For LV-mediated GABA_B_R knockout using GABA_B_1 floxed mice, animals were placed in a conditioning chamber (Context A) and, after three minutes, three conditioned stimuli (CS) (auditory tone 4 kHz, 80 dB, 30 s with 60 s interval) were presented with co-terminating unconditioned stimuli (foot shock 0.75 mA, 1 s with 90 s interval). At the end of the acquisition session, animals were removed from the conditioning chamber and returned to their home cages. Twenty-four hours later, mice were placed in a different chamber (Context B) and received only the CS without foot shock four times. Expression of fear was indexed by freezing (complete immobility) during the presentation of discrete tones and ISIs. Special care was taken to create distinct contexts by changing the shapes, flooring, ceilings, wall accessories, and odors. Recall sessions in context B were performed in almost complete darkness.

### Calculation of freezing behavior and freezing bouts

Behavioral videos were recorded from the top with a video camera at 20-30 frames per second. Mice were tracked using DeepLabCut (Mathis *et al*., 2018; Nath *et al*., 2019) with a custom-trained network for each arena and genotype (ChAT-IRES-Cre knock-in/C57BL/6J background, GABA_B_1 floxed / BALBc background) with five-point tracking (snout, left ear, right ear, center, and tail base). Tracking data was then filtered and manually inspected for quality control. To detect freezing events, we used the BehaviorDepot App (Gabriel *et al*., 2022). All acquired videos were converted to AVI format, and a velocity-based freezing classifier was then applied to the five tracked points. Each video was pre-processed by smoothing based on tracking quality and frame rate using the Lowess method (window size 12-14) and the Hampel filter (window size 6-7). The general freezing criteria for point-based tracking were a linear back velocity of less than 0.52 cm/s and an angular velocity of the head of less than 12 deg/s. Next, all immobility events were counted from 200 ms to create a histogram and identify the highest peak to determine a reliable freezing duration criterion based on the genotype. This data was then used to detect actual freezing events further. For ChAT-IRES-Cre knock-in mice, a window size of 27-32 was used based on the frame rate, with a minimum threshold count of 10 and a minimum freezing duration of 0.9 s. For GABA_B_1 floxed mice, a window size of 32 was used, based on the frame rate, with a minimum threshold count of 8 and a minimum freezing duration of 0.8 s. Using custom-written MATLAB scripts, event-related freezing time and frequency were calculated. Finally, example overlay videos of freezing events were created with custom scripts for quality checks. Total distance traveled, freezing time, frequency, and distribution were calculated using these metrics.

### Histology and immunohistochemistry

Mice were sacrificed by ketamine/xylazine overdose and perfused with 4% paraformaldehyde (PFA) in phosphate buffer (PB) for 12 minutes at a flow rate of 7.5 mL/min. Brains were then removed and placed in a 4% PFA in PB for an additional overnight at 4 degrees for post-fixation. To prevent tissue damage during sectioning, brains were cryoprotected in 30% sucrose in phosphate-buffered saline (PBS) solution overnight at 4 LJ, and 30-100 µm-thick brain slices were obtained using a sliding microtome. Slices were washed 2 times in PBS for 5 minutes each to remove any excess PFA. They were then blocked in a solution containing 10% normal goat serum and 0.1% Triton X-100 in PBS for 1 hour at RT to reduce non-specific binding. For labeling, slices were incubated with primary GABA_B_R1 antibody (Kulik *et al*., 2002) (B17 made in rabbit, affinity-purified, 0.5µg/ml in PBST 2% NGS) overnight at 4 LJ. After washing three times, slices were incubated with secondary antibody (Molecular probes, A11037, Goat anti-Rabbit IgG (H+L), Alexa Fluor 488, highly cross adsorbed, 4µg/ml) for 2 hours at RT. Slices were washed twice more in PBS for 5-10 minutes each, and in the second wash, DAPI for nuclear staining was added. Finally, the slices were mounted onto slides with a mounting medium (Mowiol). The slides were imaged using a spinning-disk (Nikon CSU-W1) or a confocal (Leica SP8, Zeiss LSM 900) microscope.

### Data analysis and statistics

For the data analysis and statistics, we used GraphPad Prism (Graphpad, San Diego, CA, USA), MATLAB (The MathWorks Inc., Natick, MA, USA), Excel (Microsoft, Redmond, WA, USA), JASP, and Clampfit (Molecular Devices) software to perform statistical tests on the data. We first checked the data for normality using the Shapiro-Wilk or Kolmogorov-Smirnov tests. For normally distributed data, we used a two-tailed Student’s t-test to compare the means of two groups. For non-normally distributed paired data, we used the Wilcoxon signed-rank test for two groups. We considered a p-value less than 0.05 to be statistically significant. We also performed repeated-measures and two-way analyses of variance (ANOVAs) with post hoc Tukey’s multiple comparisons test to compare the means of multiple groups across multiple factors. The error bars for parametric tests are presented as ± SEM. Box-and-whisker plots show the median, interquartile range (IQR), and whiskers extending to 1.5 times the IQR (Tukey method).

## Acknowledgement

We thank Bernhard Bettler for GABA_B_1 floxed mice. This project has received funding from the Austrian Science Fund (10.55776/PAT5720324) to RS, and the Marie Skłodowska-Curie grant agreement no. 665385 to CÖ. We thank Todor Asenov from the ISTA machine shop for preparing the custom part and the ISTA preclinical facility for animal care. For open access purposes, the author has applied a CC BY public copyright license to any author-accepted manuscript version arising from this submission.

---

**Supplementary figure S1. Activation of inhibitory DREADDs via CNO decreases the firing of MHb neurons in acute slices (related to Figure 6)**

**a)** Schematic representation of the experimental procedure illustrating the injection of AAV9-EF1a-DIO-hM4Di-mCherry for expressing inhibitory DREADDs (hM4Di) in cholinergic neurons of the ventral medial habenula (vMHb) in ChAT-IRES-Cre knock-in mice. The targeted pathway from the MHb to the IPN via the FR is highlighted in red. **b)** A representative image of an injected acute brain slice shows that the recorded neurons express hM4Di-mCherry (red). **c)** Representative traces of neuronal firing recorded from hM4Di-mCherry-expressing MHb neurons in acute brain slices in cell-attached mode, showing firing activity during baseline, 20 µM CNO wash-in, and the subsequent wash-out period. The trace shows a robust decrease in neuronal firing upon bath-applied CNO, indicating effective activation of inhibitory DREADDs. **d)** A representative trace of an hM4Di-mCherry-expressing neuron in the whole-cell current-clamp mode showing the basal spontaneous activity, decreased firing during the CNO application, and the subsequent recovery after the wash-out period.

**Supplementary figure S2. Intact acquisition of cued fear memory in DREADDs-injected ChAT-Cre mice (related to Figure 6)**

**a)** Graphs showing the time course of freezing behavior in 10-second bins during the acquisition phase of cued fear conditioning in mice injected with the hM4Di-mCherry-expressing AAV. Both vehicle-injected (gray, saline) and CNO-treated (orange, left; blue, right) groups show increased freezing in response to co-terminating tone-shock pairings, indicating successful fear learning. **b)** The freezing time over conditioned stimuli (CS) and inter-stimulus intervals (ISI) during the fear acquisition phase. The data depict a gradual increase in freezing with successive tone-shock pairings (CS1-ISI3), indicating similar learning curves across all groups (*main effect of stimulus: F_(3.038, 54.683)_ = 31.857, p < 0.001, repeated measures ANOVA (Greenhouse-Geisser corrected)*). These results demonstrate that DREADD-mediated inhibition in ChAT-Cre mice does not impair the acquisition of cued fear memory. Left MHb-vehicle, n = 4; Left MHb-CNO, n = 7; Right MHb-vehicle, n = 3; Right MHb-CNO, n = 8. The error bars are presented as ± SEM.

**Supplementary figure S3. GABA_B_R knock-out experimental timeline and cued fear conditioning paradigm (related to Figure 7)**

**a)** Diagram showing a timeline of the unilateral injection of Cre-expressing LV into the MHb with the pathway to the IPN via the FR, cued fear conditioning in context A on day 1, and memory recall in context B on day 2. **b)** The experimental protocol, including the acquisition (Acq., top) and recall (bottom) phases of cued fear conditioning, depicting 3 minutes of baseline habituation, conditioned stimuli (tone), unconditioned stimuli (foot shock), and the inter-trial interval with the number of repetitions.

