## Supplementary figures and images for "Asymmetrical modulation of fear expression via GABA_B_ receptors in the mouse medial habenula"

### Supplemental figure 1

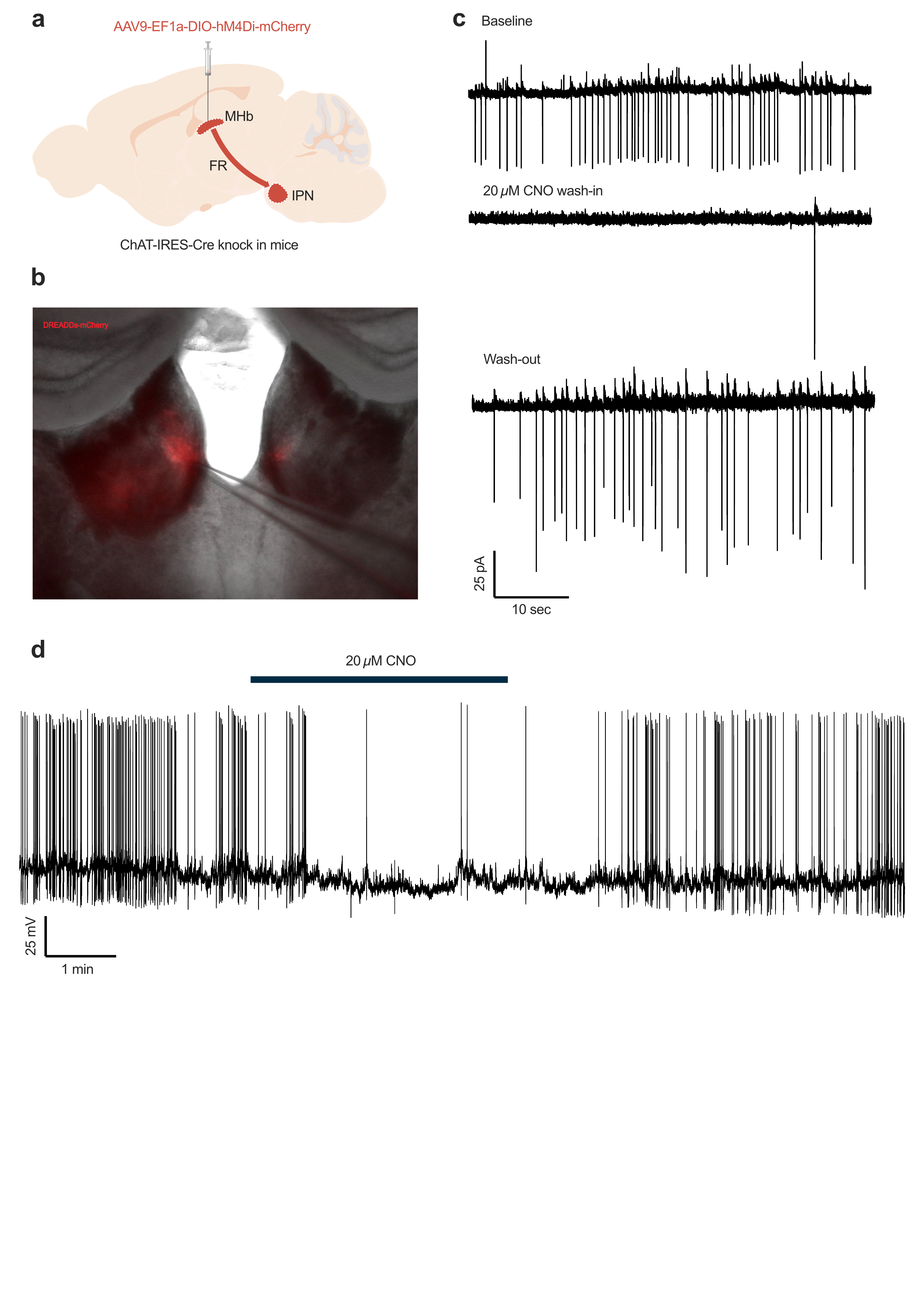

### Supplemental figure 2

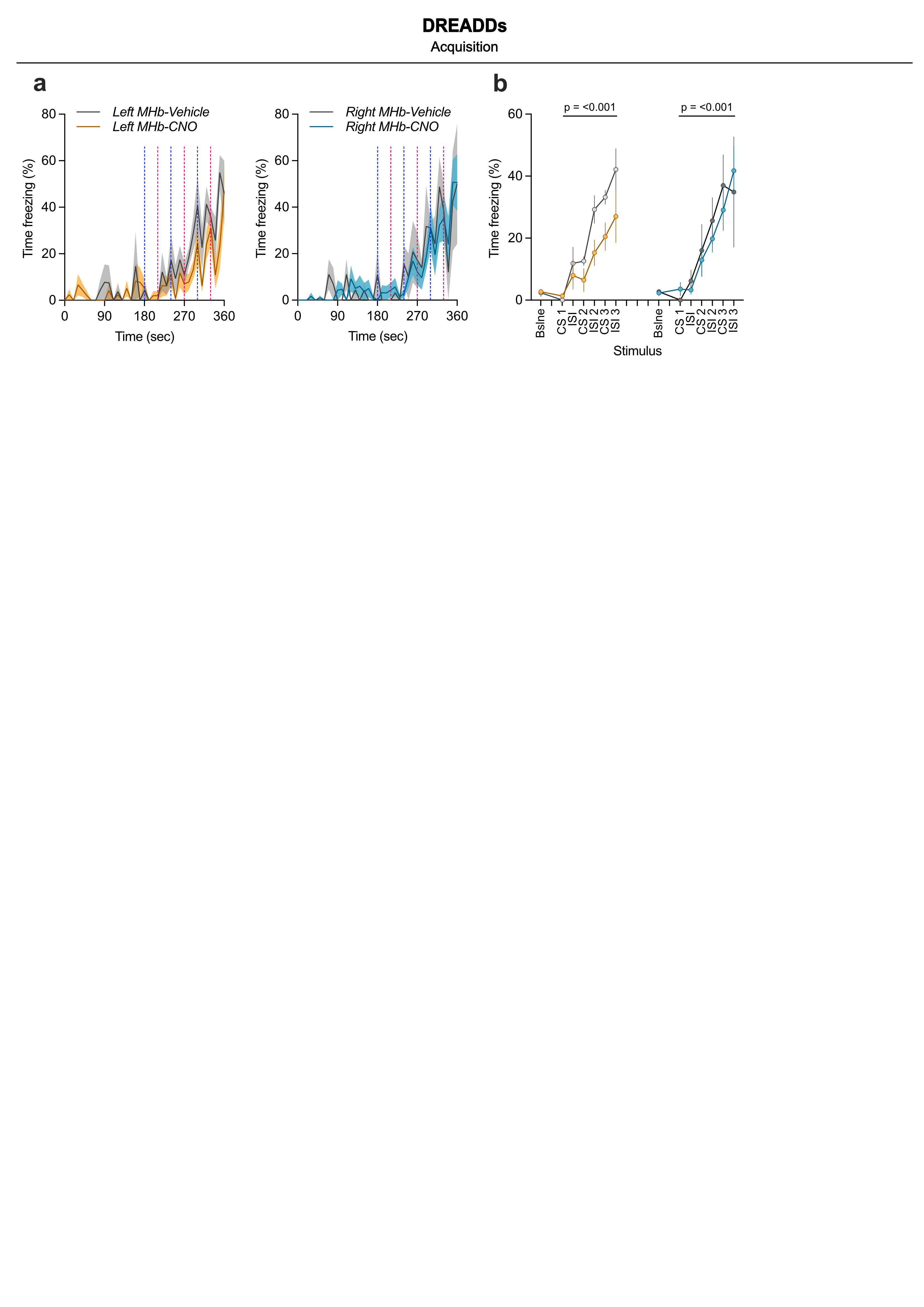

### Supplemental figure 3

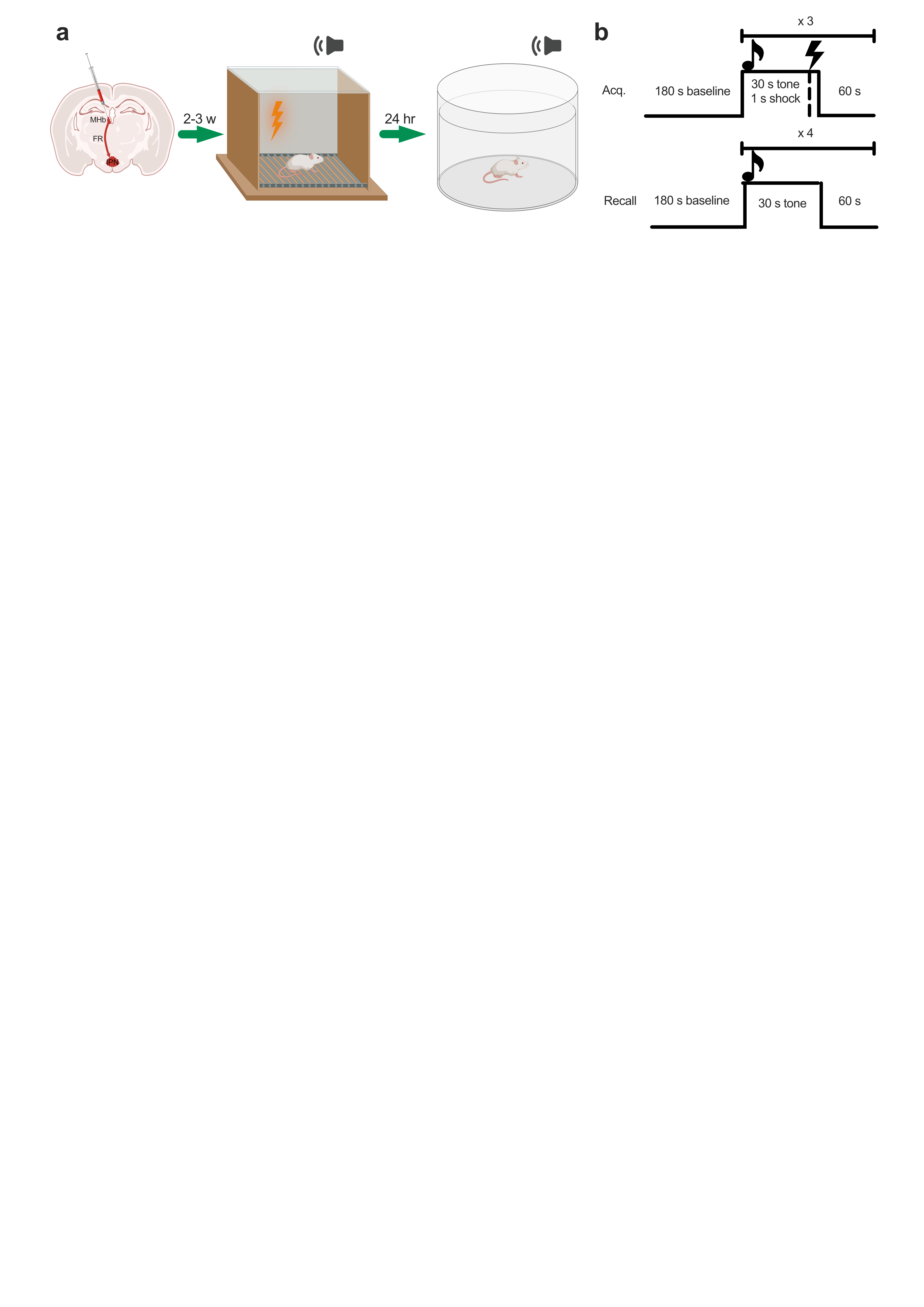
